# Sex-specific effects of appetite suppressants and stereotypy in rats

**DOI:** 10.1101/2025.02.12.637751

**Authors:** Axl Lopez, Elvi Gil-Lievana, Ranier Gutierrez

## Abstract

This study investigated the sex-specific effects of commonly prescribed appetite suppressants on body weight and the manifestation of motor side effects, specifically stereotypy. Employing video recordings and DeepLabCut (DLC) for precise behavioral quantification, we analyzed stereotypy, defined as purposeless, repetitive motor behaviors, in male and female rats. Under control (saline) conditions, male rats exhibited a greater propensity for weight gain compared to females. However, in contrast, female rats demonstrated greater and more homogenous weight loss than males following the administration of diethylpropion and tesofensine. Phentermine and mazindol induced comparable weight loss in both sexes, whereas cathine elicited weight reduction exclusively in males. 5-HTP and d-amphetamine administration only prevented weight gain relative to controls. Analysis of motor side effects revealed that drugs primarily targeting dopamine pathways – specifically, phentermine, mazindol, diethylpropion, cathine, and d-amphetamine – induced pronounced stereotypies, particularly head-weaving, in both sexes. Interestingly, tesofensine elicited head-weaving behavior exclusively in female subjects, albeit to a lesser extent than that observed with other dopaminergic agents; conversely, tesofensine was most frequently associated with orolingual dyskinesia. Moreover, one of the most potent forms of stereotypy—backward locomotion, here referred to as “moonwalking”—was sporadically observed only following the administration of phentermine, diethylpropion, cathine, and mazindol, with diethylpropion inducing it most frequently. Male subjects treated with these same drugs exhibited an unexpected effect: spontaneous ejaculations, potentially attributable to the combined effects on dopamine and serotonin signaling in brain regions regulating sexual function. Network analysis and Markov transition matrices revealed distinct behavioral profiles associated with head-weaving, which emerged as the dominant attractor state, suggesting potential mechanistic differences among these drugs. Collectively, this study provides a valuable database characterizing the behavioral side effects of appetite suppressants.

## Introduction

While Soranus of Ephesus, a Greek physician practicing in the first century AD, recognized obesity and advocated for its management through diet, exercise, and diuretics,[1] the modern obesity epidemic necessitates more sophisticated pharmacological interventions. Appetite suppressants have emerged as valuable adjuncts to lifestyle modifications, primarily functioning to reduce hunger and enhance satiety.[2] Early efforts in this domain centered on amphetamine, first employed in humans for weight reduction in the mid-20th century.[3,4] However, amphetamine’s clinical utility was hampered by a narrow therapeutic window: low doses proved ineffective for weight loss,[5] whereas higher doses elicited undesirable motor side effects, including hyperactivity and stereotypy[6,7]), besides its well established addictive properties.[8]

This limitation spurred the development of amphetamine derivatives designed to minimize adverse effects while maximizing weight loss efficacy. The prodrug diethylpropion (DEP) represented an early alternative, demonstrating some efficacy but retaining some stimulant and motor side effects [9,10]. Phentermine (PHEN), a sympathomimetic anorectic agent, is utilized as a short-term adjunct therapy for weight reduction,[11,12] but it elicits side effects such as hyperlocomotion and stereotypy.[13] Cathine (also known as D-norpseudoephedrine (NPE)), a monoamine alkaloid found naturally in the khat plant (Catha edulis) and structurally similar to d-amphetamine, has been referred to as a “natural amphetamine.”[14] Studies in rats suggest that cathine (NPE) can promote weight loss [15] and may possess a lower abuse potential compared to amphetamine.[14,16–19] Beyond amphetamines and their derivatives, other classes of appetite suppressants have been developed. Mazindol, a tricyclic imidazoisoindole non-amphetamine anorectic, has demonstrated efficacy in reducing body weight in both animal models and human trials.[20–22] Tesofensine (NS2330), a triple monoamine reuptake inhibitor and a novel investigational weight-loss agent, has shown anti-obesity effects in preclinical and clinical research.[23–25] Furthermore, research has established the role of serotonin in appetite regulation.[26] Consequently, drugs that enhance serotonergic signaling, either directly or via the administration of serotonin precursors (5-HTP), have demonstrated potential for inducing weight loss with a reduced incidence of motor side effects compared to amphetamine-based drugs.[13]

Appetite suppressants, including common amphetamine derivatives such as diethylpropion, phentermine, and cathine, are primarily employed to facilitate weight loss. These agents share a common mechanism of action, primarily involving the inhibition of serotonin, norepinephrine, and dopamine reuptake, albeit with varying affinities for each transporter, resulting in distinct pharmacological profiles. Other appetite suppressants, such as mazindol and tesofensine, are not direct amphetamine derivatives but exert similar psychostimulant effects, also acting as triple inhibitors of monoamine neurotransmitter reuptake. Additionally, the serotonin precursor 5-hydroxytryptophan (5-HTP) can exhibit appetite-suppressant properties by elevating brain serotonin (5-HT) levels. Mechanistically, appetite suppressants are designed to curb hunger and reduce food intake, promoting weight loss.[2] They exert their effects by modulating the activity of neurotransmitters in the brain that regulate appetite, including serotonin, norepinephrine, and dopamine. A common adverse effect of certain appetite suppressants, particularly those that elevate dopamine (DA) levels, is the induction of stereotypy. Stereotypy is characterized by repetitive, seemingly purposeless behaviors that lack an apparent function.[27] It can manifest in various forms, including motor instability,[28,29] head bobbing, head weaving, orolingual dyskinesias, excessive grooming, gnawing,[30] and aberrant locomotion, such as backward locomotion or “moonwalking.”[31–33]

The motor-activating effects of d-amphetamine and their derivatives in rats have been investigated not only for their central nervous system psychostimulant properties,[34,35] but also for their impact on weight loss in treated subjects.[36–38] Studies have implicated dopamine (DA) efflux from the striatum not only in the regulation of movement but also in food calorie detection, satiety, and food-related anticipatory behaviors.[39–41] While increased DA levels may contribute, in part, to the weight-loss effects of appetite suppressants, they can also precipitate motor side effects.[15,42] This is because amphetamine derivatives, by enhancing DA efflux from the striatum, influence neuronal activity in the dorsal striatum and other brain regions, such as the lateral hypothalamus and ventral tegmental area,[30] thereby promoting stereotypy and other stimulant-related side effects.[43–45] As previously mentioned, most appetite suppressants function as triple reuptake inhibitors of dopamine (DAT), serotonin (SERT), and norepinephrine (NET) transporters, indirectly elevating DA, serotonin (5-HT), and norepinephrine (NE) to supraphysiological levels, contingent upon their affinity for each transporter. Consequently, each appetite suppressant induces varying degrees of motor side effects, primarily reflecting its DA-related activity.

Despite a history spanning over half a century, a comprehensive understanding of the mechanisms by which appetite suppressants induce stereotypy remains elusive. This knowledge gap is partly attributable to the inherent difficulties in quantifying these complex and temporally extended behaviors.[46] Historically, most studies have employed subjective manual scoring to assess the magnitude of these effects,[31,33,46–49] often neglecting subtle motor kinematics over extended observation periods (hours). Similarly, ethological research traditionally relies on subjective human scoring of animal behavior, a process that is both time-intensive and susceptible to intra- and inter-observer variability. These challenges are particularly pronounced when investigating complex behaviors such as stereotypy, underscoring the need for more objective and reliable methodologies.

To address this limitation, we employed artificial intelligence (AI)-based approaches. Recent advancements in machine learning, particularly deep learning, have enabled unsupervised behavioral analysis, revealing the state of rodent behavior. One such implementation is the open-source pose estimation and movement tracking software, DeepLabCut (DLC).[50] Unlike commercially available software (e.g., Noldus Ethovision, ANY-MAZE) or other open-access video tracking tools (e.g., Bonsai), which often lack the capacity to define and score specific behaviors of interest, DLC empowers users to specify and track user-defined body parts. Consequently, DLC facilitates the accurate scoring and reporting of these behaviors, adding a crucial dimension to analyzing the animal’s movement repertoire. In this study, we utilized DLC to gain a better understanding of the motor side effects associated with certain appetite suppressants, a critical and frequently overlooked aspect of their pharmacological profile.

## Materials and Methods

### Animals

This study utilized 90 Sprague-Dawley rats (45 males and 45 females) weighing approximately 150-400 grams. To minimize stress, rats were housed in same-sex pairs and given ad libitum access to food and water. During experimental procedures, rats were individually placed in custom-designed acrylic boxes (details provided below). The housing room maintained a controlled temperature of 21°C and a 12:12-hour light-dark cycle, with lights on at 06:00 and off at 18:00.

### Drugs

The following appetite suppressants were used in this study: Phentermine hydrochloride (PHEN), Diethylpropion hydrochloride (DEP), Mazindol, D-norpseudoephedrine hydrochloride (NPE, or Cathine), Tesofensine (NS2330), 5-hydroxytryptophan (5-HTP), and Carbidopa (CB) were kindly donated by Productos Medix (Mexico). d-Amphetamine (dextroamphetamine) was kindly donated by Dr. Benjamin Floran Garduño (CINVESTAV, Mexico).

### Pharmacological administration of appetite suppressants

Phentermine, diethylpropion, mazindol, and cathine were dissolved in an isotonic saline solution. Carbidopa (CB) was suspended in a 1:4 ratio of carboxymethyl cellulose (0.5%) and saline (this mixture is referred to as “vehicle” hereafter). 5-HTP was dissolved in saline heated to approximately 40°C and kept warm until the time of injection (no longer than 20 minutes).[51] All drugs were freshly prepared and administered daily. Except for tesofensine, which was administered subcutaneously, all drugs were administered intraperitoneally. Rats were systematically weighed before the administration of either the vehicle, phentermine, diethylpropion, cathine at a dose of 20 mg/kg, mazindol, d-amphetamine at a dose of 10 mg/kg, tesofensine 2 mg/kg, and 5-HTP with carbidopa at a dose of 31 mg/kg. The doses used were based on previous reports that produce changes in body weight and motor changes, such as inducing stereotypy or hyperlocomotion.[49,10,42,52,13,21,25]

## Behavioral Procedures

### Effect of appetite suppressants on weight loss

This study investigated the effects of various appetite suppressants on the body weight of rats. Six male and six female rats were randomly assigned to one of seven treatment groups: Phentermine, diethylpropion, mazindol, cathine (NPE), tesofensine, 5-HTP, or saline (0.9%) served as a control. An additional group of three male and three female rats received d-amphetamine. All rats were housed with ad libitum access to standard rat chow (Purina, Mexico) and water. For seven consecutive days (days 1-7), rats in the appetite suppressant groups received daily injections of their assigned treatment. This was followed by a seven-day recovery period (days 8-14) without treatment. The final weight on day 14 was reported as withdrawal. The amphetamine group received daily injections but only for three consecutive days, followed by a seven-day recovery period. Body weight was measured daily 20 minutes before treatment administration between 12:00 and 16:00 hours. The change in body weight was normalized using the subject’s initial body weight at the start of the treatment, referred to as baseline (BL).

### Effect of an appetite suppressant on kinematic behaviors in an open field task

The effects of appetite suppressants on rat behavior were assessed using an open-field test. The testing arena (39 cm x 30 cm x 25 cm) was constructed of transparent acrylic and in a ventilated room isolated from external noise. Rats were habituated to the arena for two days before pharmacological treatment to minimize novelty-induced behaviors. Following habituation, each rat was individually placed into the arena, and its behavior was recorded for 4 hours using top-down and bottom-up cameras (Kayeton Technology) at a frame rate of 60 frames per second. Recordings were captured using OBS Studio software. To begin recording, the acrylic lid was placed on the arena. All tests were conducted during the inactive light phase of the rats’ light-dark cycle. Behavioral measures were recorded 15 minutes before drug administration (baseline) and 225 minutes after. Observations included stereotypy (head weaving), locomotion, quiet-awake state, rearing, and grooming. This procedure was repeated for 7 consecutive days. Kinematic behaviors were analyzed throughout the testing period using DeepLabCut, a pose estimation software that tracked six points on the rat’s body.

## Data Analyses

### DeepLabCut (DLC) Analysis

Rat behavior was analyzed using DeepLabCut (DLC) software (DeepLabCut GUI v2.2.2) [50] to track specific body points. A training dataset was created by extracting 105 frames from videos of seven rats recorded from a bottom-up view (15 frames per rat). Seven points were manually labeled on each frame: the nose, forepaws, hind paws, column (the central point between the two hind paws and the base of the tail), and base of the tail.

A ResNet-50 neural network was trained on this dataset for 1,000,000 iterations using a multi-step learning rate schedule (initial learning rate: 0.005). The trained network’s performance was evaluated by visually inspecting the labeled frames. Frames with low labeling likelihood (likelihood < 0.9) were identified and manually corrected. The network was then refined twice using the corrected labels and the same training parameters as before but with a reduced learning rate (0.001). The resulting DLC output (CSV files) containing the tracked body point coordinates was processed using custom Matlab scripts for further analysis.

### Automatic analysis of stereotypy

Analysis of head-weaving stereotypy was performed using the X and Y coordinates of the nose, hind paws, and base of the tail tracked by DeepLabCut (DLC). These coordinates were imported into a MATLAB script for further processing. To identify head-weaving events, a centroid was calculated between the two hind paw points and the base of the tail point. The angle formed by this centroid and the nose point were determined for each frame. Head-weaving stereotypy was characterized by fluctuations in this angle within a threshold of ± 2 standard deviations from the mean angle. This angular measure was further refined by incorporating locomotion data. To be classified as head-weaving, the behavior had to occur without locomotion or with only minor locomotor events lasting less than 10 seconds. Additionally, if another head-weaving event was detected within 10 seconds of the initial event, it was considered part of the same continuous head-weaving episode, even if other behaviors (locomotion, quiet-awake state, rearing, or grooming) occurred between the two events.

### Analysis of locomotion and quiet-awake state

Locomotion was analyzed using the X and Y coordinates of the forepaws and hind paws tracked by DeepLabCut (DLC). First, a centroid (center of mass) was calculated for each frame. To ensure data quality, any points with a likelihood value below 0.95 were removed and interpolated using the filloutliers function in MATLAB.

The Euclidean distance between centroids in consecutive frames was then calculated to estimate the rat’s movement. This was done using the following equation: distance = sqrt((centroid(i+1))^2 - (centroid(i))^2), where centroid(i) represents the centroid coordinates at frame i.

The resulting distance data was segmented into 600-frame snippets (corresponding to 10-second intervals at 60 frames per second) for further analysis. To classify the rat’s movement state within each snippet, the velocity and acceleration were calculated by integrating the centroid position over time.

A threshold was defined based on the maximum speed observed when the rat was in a quiet awake state. If the calculated speed within a snippet was less than or equal to this threshold, the snippet was labeled as “quiet-awake.” If the speed exceeded the threshold, it was labeled “forward locomotion.”

### Analysis of rearing and grooming

Analysis of rearing and grooming behaviors utilized the X and Y coordinates of all body points tracked by DeepLabCut (DLC). Unlike the locomotion analysis, all tracked points were included, even those with low likelihood values (likelihood ≥ 0.25). **Rearing:** A rearing event was defined by the interruption or occlusion of one or both forepaw points and the nose point from the camera’s view (likelihood < 0.80), while the hind paw points remained visible (likelihood > 0.90). This criterion reflects the rat standing on its hind legs, with its forepaws and nose raised above the field of view of the bottom-up camera. **Grooming:** Grooming behavior was characterized by rapid oscillations of one or both forepaw points (likelihood < 0.79) due to scratching movements against the face or body. This was accompanied by a high likelihood (> 0.90) for all other tracked points, indicating that the rat remained stationary during grooming. To be classified as grooming, this pattern had to be sustained for at least 100 consecutive frames (approximately 1.67 seconds at 60 frames per second).

### Automatic ethogram

Video frames were analyzed to classify rat behavior over time. Each frame was assigned to one of five mutually exclusive behavioral categories: stereotypy, quiet-awake state, locomotion, grooming, or rearing. To ensure that only sustained behaviors were captured, a minimum duration of 330 milliseconds (10 frames at 60 frames per second) was required for a behavior to be registered. While multiple behaviors could occur within a given minute, each frame was associated with only one specific behavior. This resulted in a comprehensive behavioral profile spanning the 4-hour recording period (864,000 frames). For visualization purposes, each behavior was represented by a distinct color: **Stereotypy:** Black, **Quiet-awake state:** Red, **Locomotion:** Blue, **Grooming:** Cyan, and **Rearing:** Magenta.

### Cluster classification of motor profile effects

To analyze the overall patterns of behavior across different treatment groups and time points, we performed a dimensionality reduction analysis. First, we compiled a data array where each row represented a single rat, and each column represented a specific variable. These variables included the percentage of time spent in each behavioral state (stereotypy, locomotion, quiet-awake, grooming, rearing), the onset of stereotypy, the treatment group (including both male and female groups for each appetite suppressant and the control), and the time point within the treatment period. This high-dimensional data array was then subjected to Principal Component Analysis (PCA) to reduce dimensionality while preserving the most important information. The resulting reduced-dimensionality matrix from PCA was further processed using t-distributed Stochastic Neighbor Embedding (t-SNE), a nonlinear dimensionality reduction technique well-suited for visualizing high-dimensional data, followed by a hierarchical clustering analysis, as we previously reported.[53] To determine the optimal number of clusters in the t-SNE output, we employed hierarchical clustering analysis and the elbow method.[54] This analysis suggested two distinct clusters within the data. Each point represents an individual rat, plotted according to its coordinates in the two-dimensional space defined by the t-SNE analysis.

### Markov Correlation Matrix Analysis

To analyze how the rats’ behavior changed over the treatment period, we used a Markov chain model to examine the probability of transitioning between different behavioral states. Five distinct behavioral states were defined: stereotypy, quiet-awake state, locomotion, grooming, and rearing. For each rat, we constructed a sequence of behavioral states observed over the seven days of treatment. We then identified consecutive “chains” of the same state within these sequences. For each chain, we recorded the “end state” (the last state in the chain) and the “start state” (the first state in the following chain). This allowed us to create a “transition count matrix,” where each element represented the number of times a transition occurred from a specific end state to a specific start state. We combined the transition count matrices from all rats and treatment days into a single accumulated matrix to obtain an overall picture of behavioral transitions. This matrix was then normalized by the total number of transitions observed for each end state, resulting in an “average transition probability matrix.” This matrix provided insights into the likelihood of transitioning from one behavioral state to another across the entire treatment period. This analysis was performed using a MATLAB script based on the Markov Hidden Chain algorithm,[55] explicitly utilizing the transition matrix algorithm.

### Statistical Analysis

Behavioral data were reported as mean ± SEM. Statistical differences between groups were analyzed using repeated measures ANOVA (RM ANOVA), followed by Tukey’s *post hoc* test, using GraphPad Prism 9. For a detailed description of all p values and complete statistical analysis, See **S1 Table**.

## Results

### The appetite suppressants diethylpropion and tesofensine elicited more significant weight loss in female than male rats. Phentermine and mazindol were equally effective, while cathine produced body weight reduction only in male rats

To assess whether appetite suppressants induced weight loss in a sex-specific manner, they were administered to male and female rats over 7 days, and body weight was measured after one week of drug withdrawal (**Fig 1**). The doses used were based on previous reports in which changes in body weight were observed. [10,13,21,25,42,49,56] As expected, rats treated with saline gained body weight progressively over the days in a sex-specific manner. The male rats exhibited a more pronounced weight gain than females, and these differences were significant (**Table 1**) [two-way RM ANOVA; main factor sex; F_(1,10)_ = 6.108, p = 0.0330, **Fig 1**]. One week after the final saline injection on day 14 (withdrawal), male rats weighed significantly more than female rats [t(10) = 2.911, *p* = 0.0155]. This finding confirms a tendency for male rats to gain more weight than female rats under physiological conditions. In contrast to saline, appetite suppressants induced evident weight loss or prevented weight gain, with the effects often differing between sexes.

**Fig 1.**
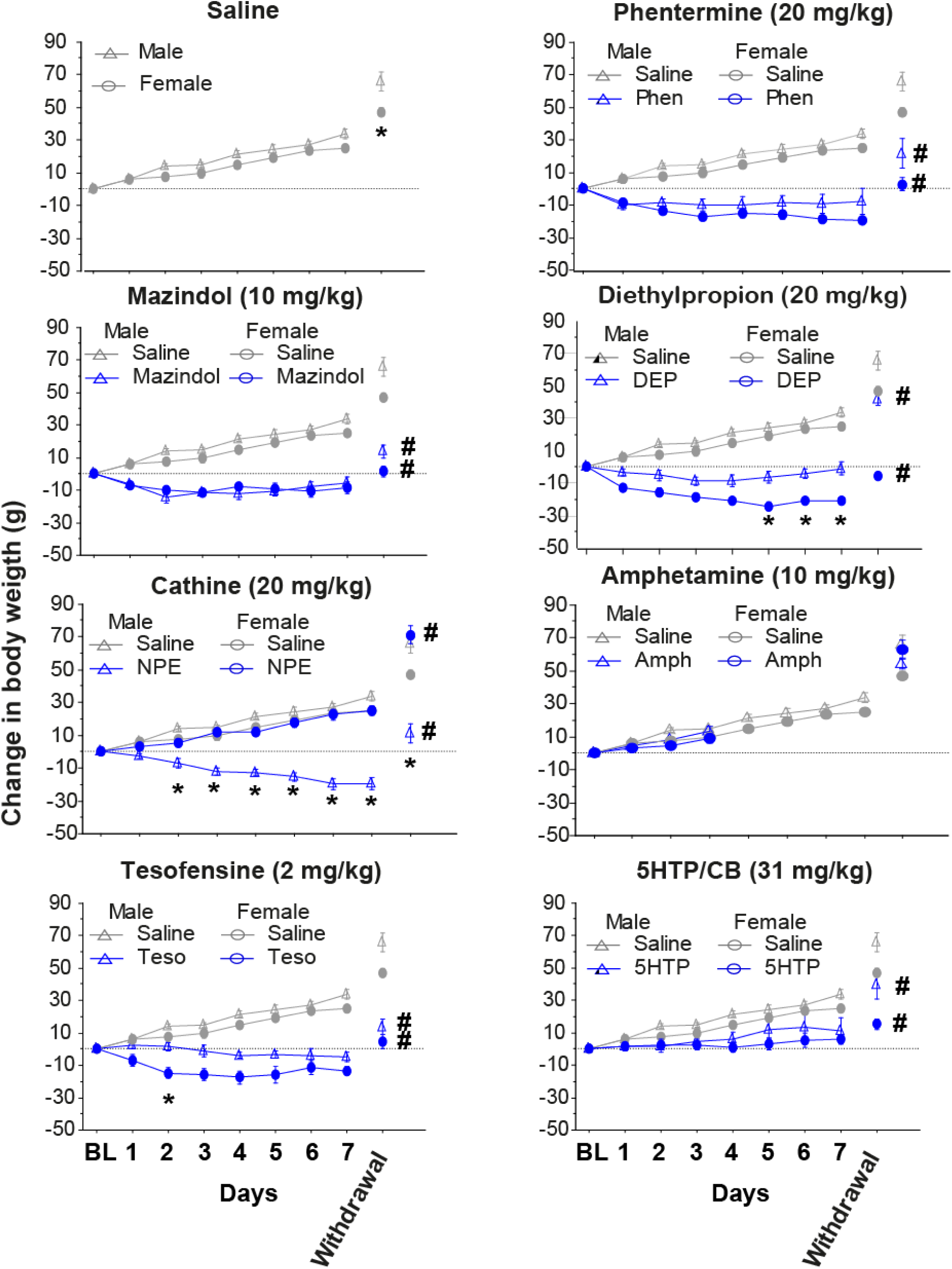
Effects of appetite suppressants on body weight change in male and female rats. The change in body weight of control naïve rats daily injected intraperitoneally with saline (n = 6 males(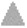 triangles), n = 6 females (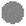 circles)) from day 1 to day 7 compared with rats injected intraperitoneally with phentermine at 20 mg/kg (n = 6 males, n = 6 females), mazindol at 10 mg/kg (n = 6 males, n = 6 females), diethylpropion at 20 mg/kg (n = 6 males, n = 6 females), cathine at 20 mg/kg (n = 6 males, n = 6 females), d-amphetamine at 10 mg/Kg (n = 3 males, n = 3 females), tesofensine (subcutaneous) at 2 mg/kg (n = 6 males, n = 6 females), and 5-HTP/CB at 31 mg/kg (n = 6 males, n = 6 females). The break in the axis indicates where the treatment was stopped. For comparison reasons, the saline group was replotted for the rest of the panels. The body weight was measured again 7 days after the last injection (withdrawal). The horizontal dotted line represents the baseline body weight before treatment. * p<0.05 for treatment days where there was a significant difference between sexes within the drug. Data is presented as mean ± SEM. # one-way ANOVA, p<0.05, significantly different from the saline group and same sex on withdrawal.

**Table 1:** Average weight change and coefficient of variation (CV) in male and female rats (for 7 days of treatment), sorted for weight loss induced in female rats.

| Drug | Mean Body Weight (F/M)(g) |  | STD Body Weight (F/M)(g) |  | CV(Female/Male)(%) |  |
| --- | --- | --- | --- | --- | --- | --- |
| Saline | 13.25 | 17.69 | 9.44 | 11.23 | 71.27 | 63.46 |
| DEP | -16.79 | -4.50 | 8.62 | 7.59 | 51.33 | 168.63 |
| Phentermine | -13.56 | -7.90 | 7.63 | 10.76 | 56.29 | 136.26 |
| Tesofensine | -11.88 | -1.60 | 9.68 | 6.63 | 81.53 | 413.25 |
| Mazindol | -7.96 | -8.56 | 6.59 | 7.86 | 82.83 | 91.74 |
| 5-HTP | 2.90 | 6.35 | 7.19 | 12.62 | 248.44 | 198.54 |
| Amphetamine | 6.63 | 9.56 | 10.10 | 14.08 | 152.46 | 147.20 |
| Cathine | 12.31 | -11.08 | 10.06 | 8.72 | 81.68 | 78.70 |

#### Phentermine

Phentermine induced a significant body weight reduction over treatment duration, reaching −13.5 g in females and −7.8 g in males, significantly different from saline control rats [three-way RM ANOVA; main factor drug: F_(1, 20)_ = 140.4, *p* < 0.0001; main factor sex: F_(1,20)_ = 5.222, p= 0.0334]. A Tukey *post hoc* unveiled no significant difference between male and female rats treated with phentermine. Importantly, this effect persisted even after one week from the last treatment (withdrawal period [one-way ANOVA: F_(3, 20)_ = 21.79 *p* < 0.0001; but no differences between sexes Tukey *post hoc* phentermine male vs. female (p = 0.1445)], with both phentermine male and female rats exhibiting significantly lower body weight compared to saline-treated rats.

#### Mazindol

Mazindol (10 mg/kg) over 7 days induced a significant weight reduction compared to saline [three-way RM ANOVA; main factor drug: F_(1, 20)_ = 184.1, *p* < 0.0001]. Like phentermine, rats administered with mazindol exhibited a similar weight reduction in females −7.9 g and −8.5 g in males (Tukey *post ho*c all p’s > 0.05), with a rapid body weight loss in the first 2 days of treatment, and then a plateau during the rest of the treatment. Interestingly, even one week after withdrawal, body weight loss remained significantly below compared to the saline group [one-way ANOVA; F_(3, 20)_ = 51.00, *p* < 0.0001].

#### Diethylpropion

Treatment with diethylpropion resulted in significant body weight reduction compared to saline treatment [three-way RM ANOVA; drug: F_(1, 20)_ = 208.5, *p* < .0001]. This effect is clearly illustrated in **Fig 1**, which shows a gradual weight loss over the first 4 days of treatment and a plateau in the last 3 days (see days 5-7). Furthermore, a significant difference in weight loss was observed between sexes [two-way RM ANOVA for diethylpropion; main factor sex: F_(1, 10)_ = 15.32, *p* = 0.0029]. Diethylpropion induced the greatest weight loss in females (−16.7 g; see **Table 1**), who lost significantly more weight than females in the control group, who gained 13.25 g. A Tukey *post hoc* analysis revealed that this sex difference was particularly pronounced during days 5-7 of treatment (*p* < 0.05), with females exhibiting a greater average weight loss (−16.7 g) compared to males (−4.5 g). This sex difference in body weight was maintained even one week after the last treatment (see **Fig 1**; Withdrawal [one-way ANOVA: F_(3, 20)_ =60.84 *p* < 0.0001; Tukey *post hoc* DEP males vs. females p < 0.0001), with females in the diethylpropion group returning to their initial baseline body weight, still significantly lower than the body weight of females in the saline control group (DEP females vs. Saline females; p < 0.0001). **Fig 1** illustrates this persistent difference in body weight during withdrawal.

#### Cathine

Unexpectedly, cathine treatment (20 mg/kg) induced opposite effects on body weight in male and female rats over the 7-day treatment period. Males exhibited an average weight loss of −11 g from their initial body weight (BL day 0), while females showed an average weight gain of +12 g. This weight gain in females was not significantly different from the +13 g weight gain observed in the saline control group. This sexually dimorphic response to cathine may be related to its interaction with sex hormones. A three-way RM ANOVA confirmed a significant interaction between sex and treatment [main factor sex: F_(1, 20)_ = 40.19, *p* < 0.0001; main factor days: F_(2.578, 51.56)_ = 31.76, *p* < 0.0001; sex x days interaction: F_(6, 120)_ = 15.77, *p* < 0.0001], indicating that the effect of cathine on weight change differed significantly between sexes (Tukey *post hoc* (days 2-7) all p’s < 0.05). This lower body weight in male than female rats persisted even during the withdrawal period [one-way ANOVA; F_(3, 20)_ = 27.52, *p* < 0.0001, Tukey *post hoc* Cathine males vs. females; p < 0.0001], whereas females significantly weigh more than saline-treated females rats (Tukey *post hoc* cathine females vs saline females p = 0.0191) as illustrated in **Fig 1**, which shows the mean body weights for each group one week after treatment cessation.

#### d-Amphetamine

A 3-day d-amphetamine administration (10 mg/kg) failed to induce significant weight loss in rats. Instead, females and males exhibited a slight, non-significant increase in body weight relative to baseline (females: +6.6 g; males: +9.5 g) comparable to the saline control group. A three-way repeated-measures ANOVA confirmed the absence of a significant drug effect [F(_1, 14)_ = 1.154, p =0.3009] or sex difference [F(_1, 14)_ = 2.727, p = 0.1209]. This lack of weight loss may be attributable to the short treatment duration or the chosen dose, despite 10 mg/kg being considered a high dose in most addiction and locomotion studies.[56]

#### Tesofensine

another appetite suppressant, also exhibited a sexually dimorphic effect on body weight loss. Female rats treated with tesofensine over 7 days exhibited a significantly greater reduction in body weight (−11.8 g) compared to males (−1.6 g), a difference of 10.2 g [two-way RM ANOVA; sex: F_(1, 10)_ = 7.295, *p* = 0.0223,] A Tukey *post hoc* revealed a significant difference between male and female rats treated in the second day of treatment(p =0.0203).This weight loss in females and males was also significantly greater than the body weight observed in the saline group [three-way RM ANOVA; main factor drug: F_(1, 20)_ = 15.77, *p* = 0.0008]. Tesofensine treatment was also efficacious in preventing weight gain (rebound) in both sexes after treatment withdrawal [one-way ANOVA: F_(3, 20)_ = 39.91, *p* < 0.0001].

#### 5-HTP/CB

5-HTP did not induce weight loss from initial baseline body weight in either female or male rats, but it significantly attenuated weight gain compared to the saline control group. A three-way RM ANOVA confirmed a significant main effect of drug treatment [F(_1, 20)_ = 15.77, *p* = 0.0008]. This effect is illustrated in **Fig 1**, which shows the mean body weight over time for each group. Importantly, significant differences between 5-HTP and saline were still observed one week after the last treatment (see **Fig 1**, withdrawal [one-way ANOVA; F_(3, 20)_ = 13.85, *p* < 0.0001]).

### Variability in body weight loss between sexes: male predisposition to weight gain under saline and enhanced female weight loss with appetite suppressant treatment

To assess sex-specific differences in the response to appetite suppressants, we examined the variability in weight change using the coefficient of variation (CV). The CV, calculated as the standard deviation divided by the mean and multiplied by 100 (CV = (SD/Mean) * 100), provides a standardized measure of dispersion relative to the mean. The CV provides insight into the consistency of weight loss responses across individuals. In the saline-treated control group, a significant sex difference in weight gain was observed. Male rats gained an average of 17.69 g, with a CV of 63%. This relatively low CV suggests that most male rats gained a similar amount of weight, indicating a consistent increase in body weight under saline treatment.

In contrast, female rats gained less weight on average (13.25 g), but their response was much more variable (CV = 71%). This higher CV reveals that some females gained significantly more or less weight than others, highlighting a less uniform response within the female group than males. This suggests an inherent difference in how males and females respond to a regular diet, with males being more prone to consistent weight gain.

When treated with appetite suppressants, the sex differences were reversed. Overall, females lost more weight and exhibited less variability (lower CVs, see **Table 1**), indicating a more potent and consistent response to the drugs. Males, on the other hand, were more resistant to weight loss and showed greater variability (higher CVs) in their response, suggesting that the effectiveness of these appetite suppressants varied considerably among individual males. This can be more clearly seen in **Table 1**.

### Appetite Suppressants Diminished Total Food Intake and Preference for High-Fat Diet (HFD) Pellets

To assess the suppression of food intake during the four-hour acrylic observation box (see below), one chow pellet (weighing between 3.6-6 g) and one HFD pellet (weighing 3.2-5.5 g) were attached to the center of one wall of the observation box each day. Pellets were placed in small plastic bottle caps secured to the wall. Food intake was measured by weighing the pellets before and after the four-hour observation period (corrected for spillage) to determine the amount consumed (see **Table 2**).

**Table 2:** Average Chow and HFD pellet intake (during 4 hours) in male and female rats (for 7 days of treatment), sorted for weight loss induced in female rats.

| Drug | HFD Intake (F/M)(g) |  | Chow Intake (F/M)(g) |  | Total Intake (F/M)(g) ± SEM |  | Preference HFD |  |
| --- | --- | --- | --- | --- | --- | --- | --- | --- |
| Saline | 4.26 | 2.44 | 0.09 | 0.79 | 4.35 ± 0.10 | 3.23 ± 0.67 | 0.98 | 0.96 |
| DEP | 0.35 | 0.23 | 0.14 | 0.08 | 0.48 ± 0.12 | 0.30 ± 0.12 | 0.71 | 0.62 |
| Phentermine | 0.22 | 0.03 | 0.12 | 0.21 | 0.34 ± 0.08 | 0.24 ± 0.11 | 0.65 | 0.19 |
| Tesofensine | 1.97 | 0.70 | 0.18 | 0.03 | 2.14 ± 0.69 | 0.72 ± 0.51 | 0.92 | 0.80 |
| Mazindol | 2.24 | 1.78 | 0.23 | 0.19 | 2.46 ± 0.58 | 1.97 ± 0.24 | 0.91 | 0.89 |
| 5-HTP | 1.00 | 0.28 | 0.95 | 0.96 | 1.26 ± 0.40 | 1.23 ± 0.28 | 0.51 | 0.23 |
| Amphetamine | 0.27 | 0.17 | 0.18 | 0.02 | 0.45 ± 0.28 | 0.18 ± 0.1 | 0.59 | 0.47 |
| Cathine | 0.81 | 0.36 | 0.17 | 0.39 | 0.98 ± 0.47 | 0.75 ± 0.40 | 0.82 | 0.67 |

Saline-treated rats, regardless of sex, exhibited a significant preference for the HFD pellet, often consuming the entire pellet. These animals consumed significantly more HFD than standard chow (**Table 2**). Conversely, diethylpropion, phentermine, d-amphetamine, and cathine significantly suppressed appetite, reducing total food intake to less than 1 g over the observation period. This marked reduction in feeding was accompanied by a decreased preference for HFD pellets, further demonstrating the efficacy of these substances as appetite suppressants.

Tesofensine (females: 2.14 g, males: 0.72 g food intake) and mazindol (females: 2.46 g, males: 1.97 g) also reduced total food intake compared to saline groups, although somewhat. Interestingly, tesofensine more strongly suppressed appetite in males than females, but males still exhibited a stronger preference for HFD pellets (preference index: males = 0.92 vs. females = 0.80). Similarly, mazindol-treated rats maintained a strong preference for HFD (preference index: males = 0.91 vs. females = 0.89). Thus, tesofensine and mazindol showed moderate effects on food intake.

In contrast, 5-HTP (females: 1.26 g, males: 1.23 g; preference index: 0.51 vs. 0.23, respectively), d-amphetamine (females: 0.45 g, males: 0.18 g; preference index: 0.59 vs. 0.47, respectively), and phentermine (females: 0.34 g, males: 0.24 g; preference index: 0.65 vs. 0.19, respectively) robustly diminished HFD intake and preference. These treatments sometimes even induced a preference for chow pellets (**Table 2**; see values below 0.5 preference index). Thus, notably, 5-HTP, d-amphetamine, and phentermine not only reduced HFD consumption but also shifted preference towards chow.

### Automatic ethogram generation from video recordings using DeepLabCut and machine learning

To evaluate motor side effects associated with each appetite suppressant, rats were video-recorded daily for 4 hours over seven consecutive days. Each rat was placed in an acrylic box and recorded simultaneously from a top and bottom view. After a daily 15-minute baseline recording period, the subject received an intraperitoneal or subcutaneous injection of either saline (vehicle) or one of the appetite suppressants (see **Methods**). These video recordings were analyzed using DeepLabCut (DLC) pose estimation software[50], combined with a machine-learning algorithm, ResNet-50, for training and refinement (see Methods for details of the model). Key points including the nose, front paws, hind paws, and base tail-were automatically labeled on the rats and remained consistent across all videos (**Fig 2A**). The extracted coordinates of these key points were used to define behaviors such as forward locomotion, rearing, grooming, and stereotypy. These behaviors were then quantitatively analyzed across all frames. From these labels, we compute all behaviors of interest (**Fig 2B**) based on the frame-by-frame coordinates of each pose estimation.

**Fig 2.**
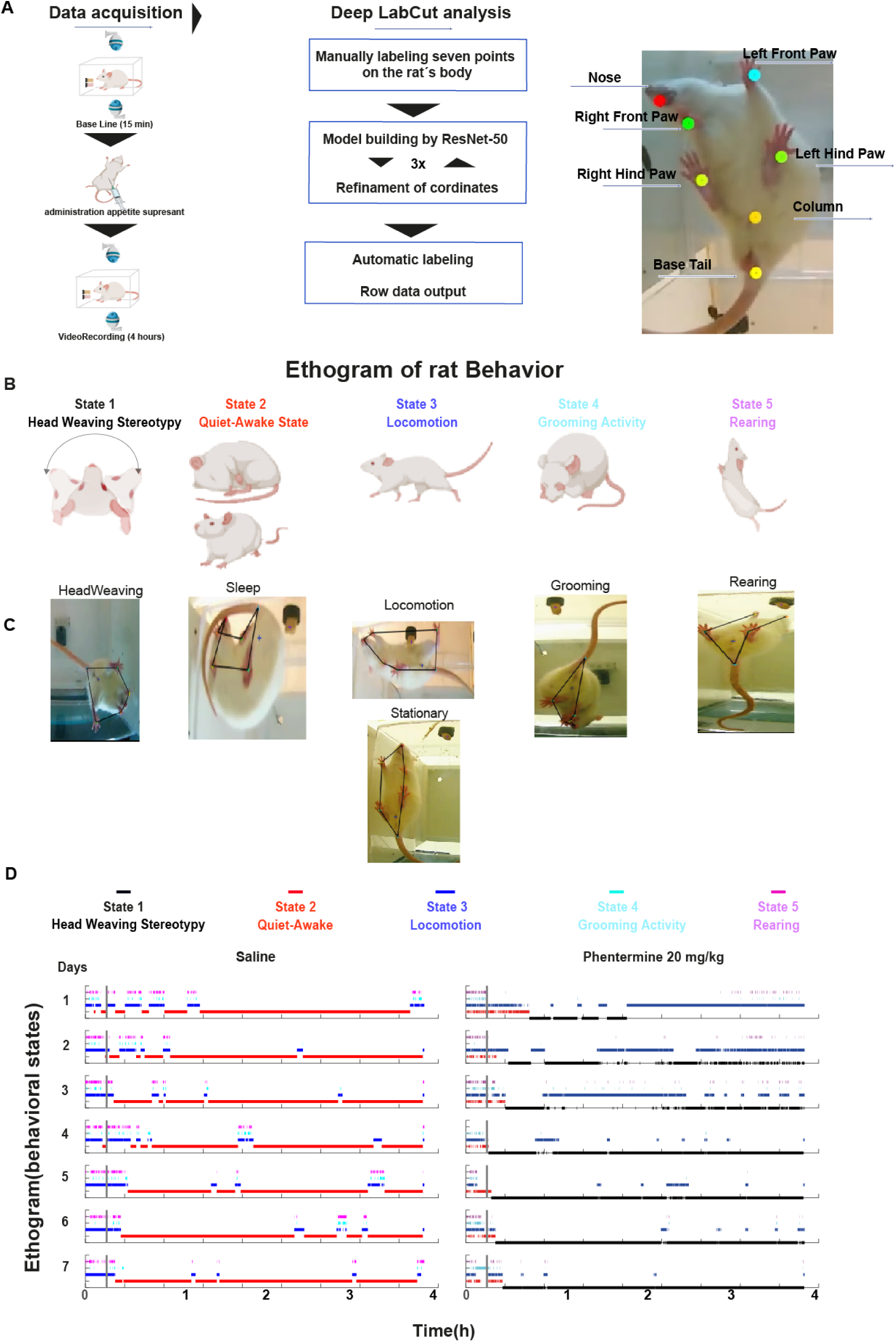
Automatic quantification of behavioral states induced by appetite suppressants. **A)** Experimental protocol for video recording rat’s behavior. For the data acquisition, rats were filmed from top and bottom views. The first 15 minutes were used as a baseline, followed by the injection of an appetite suppressant at 15 minutes. The recording continues until completing four hours. Images were converted into videos, and a machine learning algorithm, ResNet-50, was used for training these key points: the nose, front paws, hind paws, and tail base. Subsequently, the coordinates of these key points were extracted. After correcting errors, behaviors such as locomotion, quiet-awake state, rearing, grooming, and stereotypy were defined based on these coordinates and quantitatively analyzed across all frames. **B)** Ethograms were built of the following behavioral states: 1) Head weaving, 2) Quiet sleep state, 3) Locomotion, 4) Grooming, and 5) Rearing. **C)** Pose estimation images extracted from DLC show behaviors in the same order as in panel (**B**). **D)** Representative ethogram of male rats. Left panel: saline-injected rat; right panel: rat injected with Phentermine at 20 mg/kg. Behavioral states are color-coded as follows: 1) Stereotypy (black), 2) Quiet-awake state (red), 3) Locomotion (blue), 4) Grooming (cyan), and 5) Rearing (magenta). In **S1-S8 Figs,** the complete ethograms for all rats and drugs can be seen, and the entire database can be found in [87].

From DLC-tracked points, we formed polygons by connecting points such as nose-front paws (top point), left front paw-left hind paw-base of the tail, right front paw-right hind paw-base of the tail, and left hind paw-base of the tail-right hind paw (bottom view) (**Fig 2C**). These polygons allowed us to build the ethograms frame by frame, indicating whether an animal was in a state of stereotypy (head weaving), quiet-awake state (including putative sleep periods and inactivity), locomotion, grooming, or rearing (**Fig 2D**). To calculate the stereotypy behavior, we used the centroid formed by the joints between the front paws and the nose, measuring the angle difference (between frames) relative to the one formed by the hind paws and the base of the tail (see Methods). For rearing and grooming behaviors, the tracking of these polygons helped identify moments when the edges between the front and hind paws deformed or disappeared, providing a key indicator for determining these behaviors (see **Methods**).

Analysis using the automatic ethogram revealed that saline-treated rats (**Fig 2D**, left panel) did not exhibit any head-weaving stereotypy. Instead, they spent most of the observation period in a quiet-awake state. Initially, these rats displayed exploratory behavior, with long periods of forward locomotion alternating with rearing. After one hour, this exploratory behavior decreased almost entirely. Following this, the animals entered prolonged periods of inactivity, characterized by a quiet-awake state (standing motionless) or sleep postures such as curling up in a ball. This transition occurred more rapidly across days, decreasing from an average of ∼60 minutes on day 1 to ~20 minutes on day 7 as the rats acclimated to the observation box.

In contrast, rats treated with some appetite suppressants, such as phentermine, exhibited no sleep or inactivity after injection. Instead, they displayed pronounced head-weaving stereotypy (see **Video 1** for males and **Video 2** for females), spending approximately >50% of the observation period in this purposeless behavior. Moreover, the onset of head-weaving became progressively faster over the days of treatment, with the latency to onset decreasing from an average of 56.50土19.67 SEM minutes on day 1 to 7.46土1.59 SEM minutes (post-injection) on day 7 (**Fig 2D**, right panel), reflecting a sensitization process. In subsequent days, head-weaving became the predominant activity, with a strong tendency for the animals to remain in this state once it is initiated. **Fig 2D** (right panel) depicts these behavioral patterns by showing a representative ethogram or the time course of behavioral states of one rat across seven days.

### Control saline-treated rats

Once the motor behaviors associated with each appetite suppressant were computed (**Fig 2D**), we calculated the total time spent on each behavior over the 4 hours of recordings. Vehicle-treated rats remained predominantly in a quiet-awake state (including sleep-like behavior) most of the time, slightly increasing throughout the treatment (**Figs 3A males** and **4A females**). This could be due to the rats becoming more familiar with the behavioral box over time (See **S1 Fig** for all ethograms of saline-administered rats).

**Fig 3.**
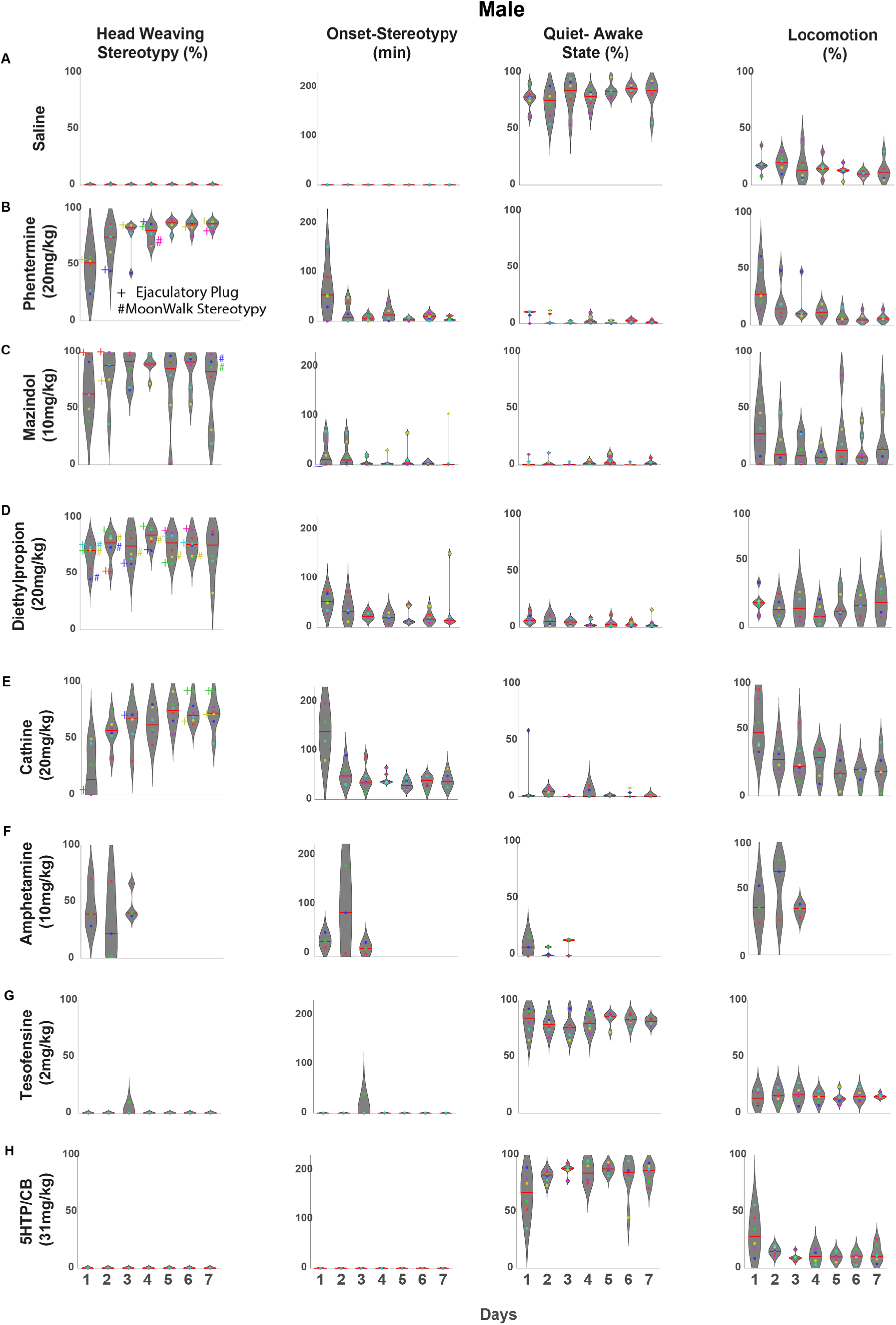
Effect of appetite suppressants on head-weaving stereotypy in male rats. Drugs were ranked in descending order of potency on stereotypy. Each behavior is quantified as a proportion of the time spent in that state during the 4 h duration of each session. Each dot within the violin plots represents an individual rat, and the colored dot corresponds to the same rat across various days and behaviors. **A)** Saline: As expected, under physiological conditions, rats spend most of their daytime in a quiet-awake time (immobile but mainly in a sleeping-like posture) or exploring their environment (locomotion). No head-weaving stereotypy was observed. **B)** Phentermine (20 mg/kg): Stereotypy strengthens over the days, occupying more than 60% of session time from the fourth day onwards. Moreover, we observe a progressively shorter onset of stereotypy across days. **C)** Mazindol (10 mg/kg) also induced significant head weaving stereotypy, although with more variability than phentermine. **D)** Diethylpropion (20 mg/kg) was the third appetite suppressant with a robust stereotypy, although its stereotypy was more homogeneous than phentermine (CV for DEP = 17.85% vs CV for PHEN = 22.01%). **E)** Cathine 20 mg/kg (d-Norpseudoephedrine (NPE)): Stereotypy was not as pronounced as with phentermine, and rats exhibited high variability (CV = 37.81%), but it tended to strengthen across days. **F)** d-amphetamine (10 mg/kg): High but variable levels of stereotypy were observed from the first day of treatment (CV =53.35%). Additionally, this drug induced a quiet-awake state (including sleep-like behaviors) on the first and third days of treatment, suggesting that their effects disappeared more rapidly than other appetite suppressants. **G)** Tesofensine (2 mg/kg): Unlike other appetite suppressants, males do not exhibit significant head weaving stereotypy, as previously reported [25], except for one subject on the third day. Tesofensine-treated rats exhibited prolonged periods of immobility (quite-awake state) while remaining awake, alternating with brief sleep-like posture, suggesting this drug may have insomnia-inducing properties. **H)** 5-HTP (31 mg/kg): 5-HTP/CB treatment did not induce head-weaving stereotypy. Rats treated with 5-HTP exhibited behavioral profiles similar to saline controls, characterized by prolonged periods of inactivity, primarily sleep-like behavior. For all the drugs, the symbols # and + and the color next to each dot indicate the rat and the day on which a subject presented spontaneous ejaculation (+) and/or backward locomotion (# moonwalk stereotypy)

### Dopamine-targeting drugs induced head-weaving stereotypy in both sexes

The following section details the motor side effects of appetite suppressants, ranking drugs from most to least severe.

#### Phentermine

In contrast, phentermine (**Fig 3B males**) predominantly induced head weaving stereotypy, which became more pronounced over the days [one-way ANOVA F_(6,35)_ = 6.865, p < 0.0001] a Tukey *post hoc* revealed that the first day was significantly different from the third day onwards (Tukey *post ho*c all p’s < 0.05) not only in duration but also in having a faster onset [one-way ANOVA F_(6,35)_ = 5.220, p = 0.0006]. A Tukey *post hoc* revealed a significant difference between the first day versus the rest of the days (All p’s < 0.05). While some subjects treated with phentermine exhibited high levels of locomotion during the first two days of treatment, locomotion gradually decreased over the days. This decrease in locomotion coincided with the opposite increase in stereotypy. Therefore, although initial locomotion was high in some phentermine-treated subjects, this effect was likely offset by the later increase in stereotypy, resulting in no overall difference in locomotion compared to the vehicle group [two-way RM ANOVA; main factor; drug: F_(1, 10)_ = 0.000291, *p* = 0.9867]. Moreover, we observed an inverse correlation between the increase in the percentage of time spent in stereotypy and a decrease in locomotion over days in males treated with phentermine (Pearson correlation coefficient, r = −0.9941, p < 0.0001).

Similarly, female rats (**Fig 4B**) treated with phentermine predominantly exhibited head-weaving stereotypy but in a more homogeneous manner than males from the first day (males had a larger CV= 22.28% vs. females CV=10.54%), as all female subjects spent more than 50% of the time in a head-weaving state from day one and subsequent days (**S2 Fig**). In females, there is also an effect where levels of stereotypy increase over the days, whereas locomotion decreases (Pearson correlation coefficient, r = −0.8364, p < 0.0001). These findings suggest that, throughout the treatment, the predominant side effect of these drugs shifts toward head weaving stereotypy, gradually taking over locomotor activity.

**Fig 4.**
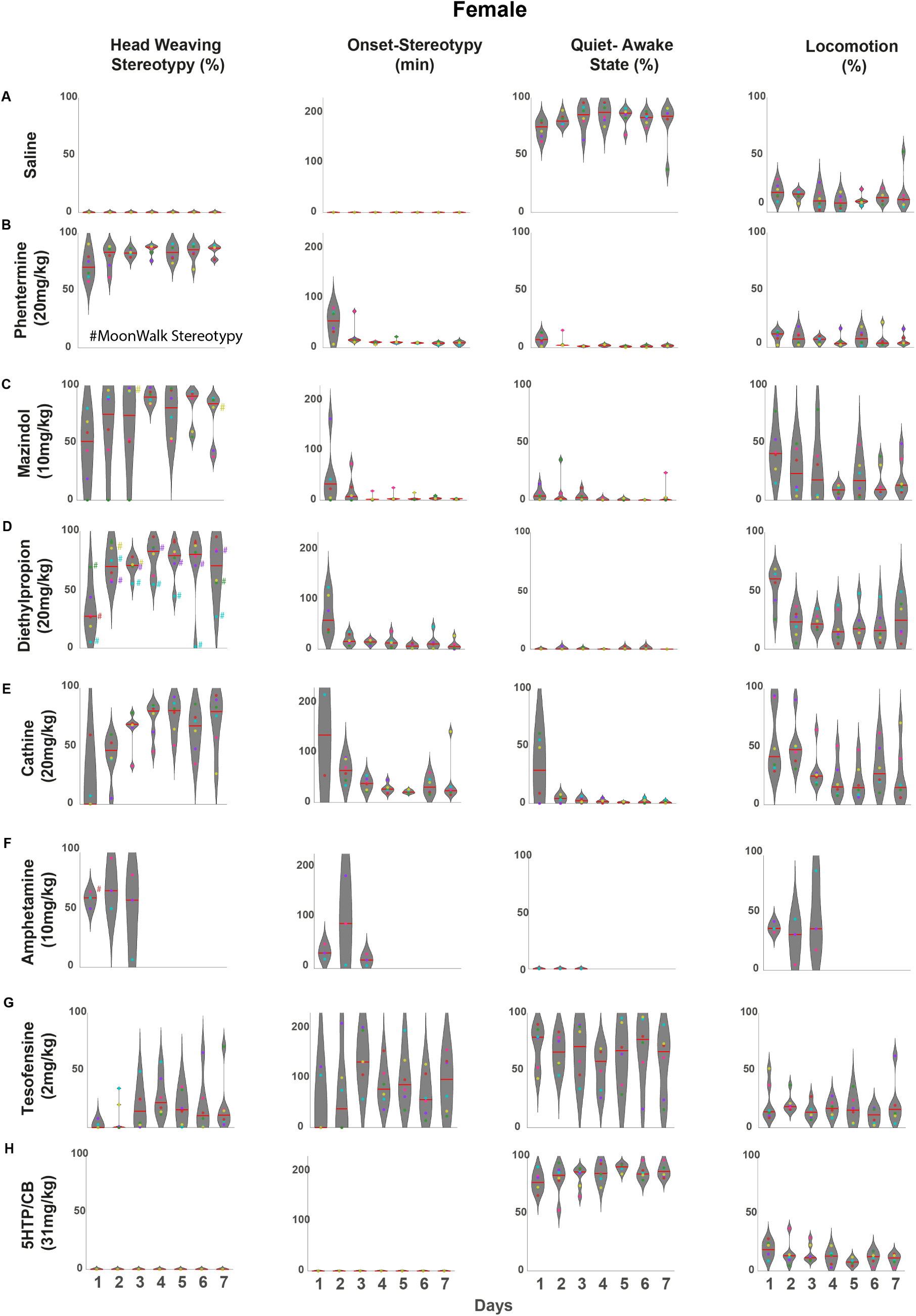
Effect of appetite suppressants on head-weaving stereotypy in female rats. Drugs were ranked in descending order of potency on stereotypy. Same conventions as in Fig 3, but for female rats: **A)** Saline: female rats spend most of their time in the quiet-awake state (mostly sleeping). **B)** Phentermine (20 mg/kg): Induced stereotypy in females is even more homogeneous than in males from the first day (males had a larger coefficient of variation in treatment CV = 22.2% versus females CV = 10.5%. Moreover, the onsets are quicker from the second day of treatment, with rats exhibiting head-weaving behavior within 15 minutes post-injection. **C)** Mazindol (10 mg/kg): Similar to males, female rats exhibit marked head weaving stereotypy. However, one subject (green dot) stands out, as it exhibited very low levels of stereotypy but high locomotion during the first three days of treatment. **D)** Diethylpropion (20 mg/kg): In contrast to the low variability shown by males (CV = 17.8%), females exhibited high variability in the induced levels of stereotypy (CV = 35.9%). A particular case stands out with the cyan dot, which consistently showed stereotypy levels below the mean except for the second day of the treatment. **E)** Cathine 20 mg/Kg (d-Norpseudoephedrine NPE): Cathine was the drug that induced the fourth-highest average level of stereotypy in both females and males. However, unlike males (CV = 37.8%), there was significant variability between subjects (CV = 49%) and across days. For example, the subjects represented by the red dot showed increasing levels of stereotypy over the days, whereas the subjects represented by the cyan dot transitioned from high levels of stereotypy during the first days of treatment to levels below the mean on days 6 and 7 of treatment. **F)** d-amphetamine (10 mg/kg): From the first day of treatment, subjects exhibited a level above 50% of time spent on stereotypy, which slightly increased on the second day. Notably, one subject (cyan dot) showed a considerable decrease in stereotypy levels on the third day, quantified after 100 minutes post-injection. **G)** Tesofensine (2 mg/kg): In females, tesofensine, unlike in males, did induced head weaving behavior, although with high variability between subjects (CV = 116.04%), which was also characterized by its delayed onset (82.62 min), highlighting a gender-specific effect of the drug. **H)** 5-HTP (31 mg/kg): Like in male rats, serotonin alone does not induce the head-weaving stereotypy. For all the drugs, the symbols # and color next to each dot indicate the rat and the day a subject presented backward locomotion (# moonwalk stereotypy).

#### Mazindol

Mazindol induced the second most potent effect on stereotypy. **Figs 3C** (male) and **4C** (female) show that chronic administration of mazindol led to head-weaving stereotypy in both sexes on each treatment day. However, rats exhibited a large variability. For example, two male rats exhibited low levels of stereotypy (see **Fig 3C**, yellow and cyan dots, **S3 Fig** see male rats 4 and 6) but high locomotion. Likewise, one female rat had no stereotypy on the first three days (**Fig 4C** green dot, **S3 Fig,** see female rat 5), but on days 4, 5, and 7, exhibited more than 90% of the time stereotypy. Another female rat (**Fig 4C** pink dot) spent less than 50% in stereotypy on most days except day 6 (>80%), but they spent extended periods in rearing (see **S3 Table**, **S6 Fig**). This variability was reflected in a high CV = 40.04%. Also in both, male and female rats exhibited an anti-correlation between the stereotypy and locomotion behavior, where levels of stereotypy increase over the days, whereas locomotion decreases (Pearson correlation coefficient for males, r = −0.9878, p < 0.0001, and for females r = −0.9641, p < 0.0001).

#### Diethylpropion

In male rats (**Fig 3D**), diethylpropion, like phentermine and mazindol, was the third drug that induced a stronger head-weaving stereotypy in all subjects throughout the 7 days of treatment. Stereotypy gradually increased over the days, reaching its peak level of time spent in this behavior (82.7% of the 4-hour observation time was spent on stereotypy) on the fourth day of treatment, after which it plateaued during the last three days. A strong anticorrelation was observed between the increase in stereotypy and the decrease in locomotion (Pearson correlation coefficient, r = −0.9194, p < 0.0001).

Diethylpropion induced head-weaving stereotypy in both sexes, but with a significant difference in the magnitude of the effect (See **S4 Fig** for all ethograms). On the first day of administration, males exhibited head-weaving stereotypy 65.93% of the time, markedly higher than the 32.52% observed in females (**Fig 4D**). Conversely, females showed significantly greater locomotor activity (54.43%) than males (22.99%). A strong negative correlation confirmed this inverse relationship between stereotypy and locomotion (Pearson correlation coefficient, r = −0.9450, p < 0.0001).

#### Cathine

Cathine also proved to be an effective appetite suppressant in male rats, though it notably induced head-weaving stereotypy, making it the fourth drug identified in this study to elicit this behavior. On the first day of treatment, male subjects (see **Fig 3E**, blue and cyan dots) showed levels of stereotypy under the mean in all of the seven days of treatment, and this made cathine the highest variability (CV = 37.81%) in males among the appetite suppressants that successfully induced stereotypy in all subjects. However, the same as in phentermine, the levels of stereotypy gradually increased over the days, reaching their peak by the fourth and fifth days of treatment.

In females, cathine showed diverse stereotypy patterns, with some subjects presenting very high levels of stereotypy (see **Fig 4E**, green and yellow dots), while other subjects displayed predominantly locomotion rather than stereotypy (see **Fig 4E** violet dot and **S5 Fig**). A phenomenon observed in both sexes was that stereotypy levels were low or absent on the first day of treatment, gradually increasing over time. However, females exhibited significantly lower stereotypy levels on the first day of treatment (11.4%) compared to males (20.60%), highlighting a sex-dependent difference in the initial response to cathine.

#### d-Amphetamine

We administered chronically for three days in both male and female rats a high dose of 10 mg/kg (given that the lethal dose is in the range of 10-30 mg/kg, and thus we decided to treat rats for only 3 days)[57]. It has been reported that doses starting at 2.5 mg/kg are enough to induce locomotion in rats,[52,58] while doses higher than 5 mg/kg start to induce stereotypy.[49,52,59] Beyond 10 mg/kg, the stereotypes do not differ much, so we used a dose of 10 mg/kg to analyze stereotypy head weaving. In male rats, d-amphetamine induced head-weaving stereotypy with a rapid onset on the first day (average onset time: 49 minutes). However, the percentage of time spent in this state was relatively low (46%) due to a significant amount of drug-induced locomotion (44%), which increased further on the second day of treatment (63% of the observation time). Despite the strong inverse correlation between stereotypy and locomotion (Pearson correlation coefficient, r = −0.94402, p = 0.00001) (**Fig 3F**), d-amphetamine exhibited a unique behavioral profile. In contrast to other appetite suppressants, where stereotypy predominates, d-amphetamine-induced both increased locomotion and decreased stereotypy. Visual observation revealed more frequent head movements during locomotion than saline-treated rats (**S3 Video**).

While d-amphetamine elicited high levels of stereotypy with an early onset on the second and third days of treatment, some subjects transitioned into a quiet-awake state, including a curled-up posture indicative of a sleep-like state, towards the end of the observation period (see **Fig 3F**, blue and green dots, and **S6 Fig,** see male rats 1 and 2). This shift from stereotypy to sleep-like behavior in the latter half of the observation period was unique to d-amphetamine among the dopaminergic appetite suppressants tested, potentially suggesting rapid clearance at this dose.

In females, rats exhibited stereotypy throughout the three days of treatment (**Fig 4F**). Some subjects exhibited low levels of stereotypy (see **Fig 4F**, purple and cyan dots) but high levels of locomotion (**S6 Fig** see rats 2 and 3).

### Sex-dependent behavioral effects of tesofensine: Greater susceptibility stereotypy in female rats

A previous study from our group reported that tesofensine 2 mg/Kg in male rats failed to induce head weaving stereotypy, which was only present at a high dose of 6 mg/Kg accompanied by orolingual dyskinesia.[25] However, we did not explore the effect of tesofensine in female rats. Here, we investigated whether the appetite-suppressant tesofensine induces stereotypy in both sexes. Male rats administered 2 mg/kg tesofensine spent significantly more time in a quiet-awake state (including curled-up in a ball posture, a sleep-like posture, a difference notably more pronounced than in female rats [RM ANOVA; sex factor: F_(1,10)_ = 16.31, p = 0.0024]. Only one male exhibited brief stereotypic head-weaving on the third day of treatment (**Fig 3G** green dot and **S7 Fig** see rat 2). In contrast, females showed significantly more pronounced stereotypy than males across sessions [RM ANOVA; sex factor: F_(1,10)_ = 50.17, p < 0.0001]. Interestingly, in most cases, tesofensine-induced stereotypy appeared later (in the last hour) in the recording session and was characterized by subtler head movements compared to the more intense head-weaving behaviors induced by other appetite suppressants (**S7 Fig**). These findings suggest that female rats are more susceptible than males to tesofensine-induced weight loss (**Fig 1**) and behavioral motor side effects.

### 5-HTP, a serotoninergic precursor, did not induce head-weaving stereotypy

To evaluate the chronic effects of 5-HTP/carbidopa (CB) on stereotypy expression, male and female rats received daily administrations of 31 mg/kg 5-HTP combined with 75 mg/kg CB, a peripheral inhibitor of aromatic L-amino acid decarboxylase that indirectly elevates brain serotonin (5-HT) levels [60]. This regimen did not elicit measurable head-weaving stereotypy in either sex during the four-hour observation period (**Figs 3H, 4H**, and **S8 Fig**). Furthermore, the behavioral profiles observed following 5-HTP/CB treatment were largely indistinguishable from those exhibited by the vehicle group (e.g., compare **Figs 3A and 3H**).

### t-SNE and hierarchical clustering analysis revealed two distinct motor behavioral profiles induced by appetite suppressants

To further explore the overall map of motor effects of appetite suppressants, we investigated whether these drugs induced distinct clusters of motor behaviors. We used t-SNE, a dimensional reduction technique, and hierarchical clustering analysis to map all motor effects of each rat into clusters based on shared behavioral characteristics (stereotypy, locomotion, quiet-awake state, grooming, rearing), where each point represents an individual rat (**Fig 5**). This analysis revealed two primary groups: 1) rats treated with 5-HTP/CB and male rats administered tesofensine (2 mg/kg), which exhibited motor profiles similar to saline-treated controls, and 2) rats treated with dopaminergic drugs (phentermine, mazindol, diethylpropion, and cathine), all of which induced substantial stereotypy, formed a distinctive cluster (**Fig 5** see arrow stereotypy cluster). Interestingly, tesofensine exhibited unique sex-dependent dynamics. Female rats treated with tesofensine (2 mg/kg) formed a smaller, intermediate position between clusters, suggesting a motor profile with features shared between control-like behaviors and stereotypy-inducing drugs. Additionally, three outlier rats treated with cathine (two males and one female) displayed unique motor profiles that did not fit into the main two clusters. Together, these findings illustrate that drugs that elevate synaptic dopamine levels evoke a markedly distinct motor behavior profile compared to drugs with little to no effect on dopamine reuptake inhibition.

**Fig 5.**
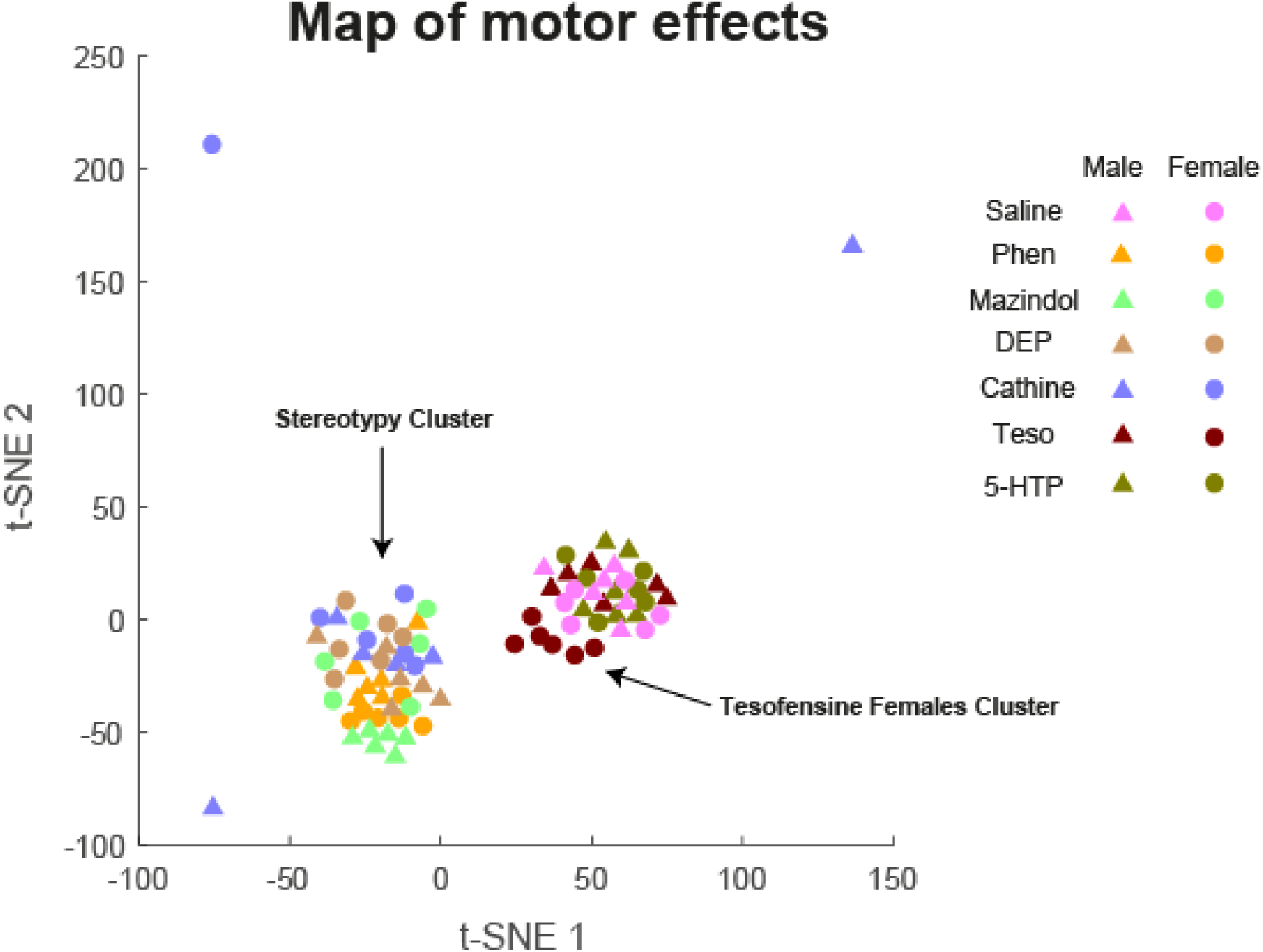
t-distributed Stochastic Neighbor Embedding (t-SNE) and hierarchical clustering analysis delineate distinct clusters of drug-induced motor effects in rats, with tesofensine in females demonstrating an intermediate profile. The analysis reveals two primary clusters: one encompassing amphetamine-like drugs associated with stereotyped behaviors and another comprising drugs that do not elicit such behaviors. t-SNE map illustrating the clustering of motor effects across various treatments. Each data point represents an individual subject, with triangles indicating males and circles indicating females. Colors denote different treatments: saline (pink), phentermine (yellow), mazindol (light green), diethylpropion (beige), cathine (blue), tesofensine (burgundy), and 5-HTP (dark green). Tesofensine-treated females (burgundy circles) are positioned intermediately between the two main clusters. This suggests that tesofensine in females elicits a motor effect profile that shares characteristics with both amphetamine-like and non-stereotypy-inducing drugs.

### Behavioral state transitions induced by appetite suppressants

To determine whether transitions between behavioral states varied by drug, sex, or an interaction between both, we constructed a weighted network using Markov chain transition probabilities between behavioral states.[55] This approach allowed us to compare the predominant transition pattern in response to each drug. In saline-treated rats, both sexes exhibited highly similar transition dynamics between states (**Fig 6B**), with the quiet-awake state being the most prominent (red node, the diameter of each circle is proportional to the fraction of time the rat spent engaged in that behavior). These rats transitioned primarily from quiet-awake to locomotion, followed by rearing, revealing the most common physiological transitions between behaviors. Impressively, both transition networks were highly similar in male and female rats.

Conversely, phentermine, mazindol, diethylpropion, and cathine produced a more active pattern (**Figs 6C, 6D, 6E**, and **6F**), where head-weaving stereotypy emerged as the primary state (see the large diameter of the black node), commonly followed by transitions from stereotypy to locomotion. This network analysis also revealed a similar transition frequency between male and female rats treated with these appetite suppressants (see **Figs 6C-F**).

In contrast, d-amphetamine and tesofensine (**Figs 6G** and **6H**) displayed unique sex-dependent dynamics: d-Amphetamine also induced distinct motor patterns (**Fig 6G**). Male rats transitioned from locomotion (blue node) to head-weaving (black node), rearing (light blue node), and a quiet-awake state (red node). In contrast, female rats more frequently transitioned between locomotion (blue node) and head-weaving (black node).

Likewise, tesofensine-treated males transitioned predominantly from locomotion (blue node) to grooming (pink node, **Fig 6H**) (also see the tiny black node of stereotypy), while females exhibited a main transition from locomotion (blue node) to head-weaving stereotypy (black node).

### Other forms of stereotypy: orolingual dyskinesia and moonwalk locomotion

In addition to head-weaving stereotypy, another stereotypy observed includes orolingual dyskinesias (tongue protrusion and retraction in the air, **S4 Video**). This dyskinesia was particularly strong after the administration of mazindol. Tesofensine induced sporadic, brief, and abnormal jaw and tongue movements (orolingual dyskinesia) in both male and female rats during periods of quiet wakefulness (**S4 Video**). Notably, this effect was observed in male rats despite the absence of head-weaving stereotypy in males (**Fig 3G**, also see **S4 Video**), suggesting that tesofensine may induce distinct motor effects compared to other drugs. Previously, we reported that a high dose of 6 mg/Kg of tesofensine induced robust orolingual dyskinesia in male rats.[25]

Moreover, male and female rats treated with phentermine, diethylpropion, cathine, d-amphetamine, and mazindol exhibited additional stereotypes. For example, these drugs led to a unique gait pattern here, referred to for the first time as “moonwalk” locomotion. The moonwalk stereotypy was exclusively seen in males when using phentermine (**S5 Video, Fig 3B**), while it occurred more frequently in both males and females with diethylpropion (**Figs 3D** and **4D**; **S6 Video and S7 Video**), mazindol (male **Figs 3C** and female **4C**; see **S8 Video** and **S9 Video**). In the case of d-amphetamine, this backward locomotion was only observed in females (**Fig 4F** see # symbol, **S10 Video**). A systematic and formal measurement of orolingual dyskinesia and moonwalk effects is out of the scope of this manuscript. However, these behaviors further underscore the potent dopaminergic effects of the drugs, reflecting their ability to induce a broad spectrum of repetitive and abnormal purposeless motor patterns.

### Appetite suppressants and spontaneous ejaculations

Spontaneous ejaculations (**Fig 3C** see + symbol) were observed in male rats treated with appetite suppressant phentermine (**S11 Video**), mazindol (**S12 Video**), diethylpropion (**S13 Video**), and cathine (**S14 Video**). This side effect, which wasn’t seen with other drugs such as d-amphetamine (mainly DA releaser and blocker of transporter DAT,[61] 5-HTP (a pure serotoninergic precursor), highlights that the pronounced dopaminergic or serotonergic activity per se of these compounds is not sufficient to induce spontaneous ejaculation and semen discharge. Therefore, spontaneous ejaculations may be due to the drugs’ effects on combined effects on dopamine and serotonin signaling in brain regions involved in sexual function.[62–64] This is because a synergistic effect between DA and 5-HT has been reported on semen discharge.[64–66]

## Discussion

This study examined the effects of several appetite suppressants on weight loss and motor behaviors in both male and female rats. In general, female rats demonstrated more consistent weight loss, exhibiting reduced inter-individual variability compared to males. Specifically, female rats exhibited significantly greater weight loss than males in response to diethylpropion and tesofensine. Conversely, and unexpectedly, cathine induced weight loss only in male rats. In terms of motor side effects, tesofensine generally induced less pronounced stereotypy than the other compounds tested. However, drugs primarily targeting dopamine pathways – namely, phentermine, diethylpropion, mazindol, and cathine – elicited marked stereotypies, particularly head-weaving (male **S1 Video** and female **S2 Video**). Notably, tesofensine also induced head-weaving behavior, specifically in female rats, although this effect was not as pronounced as with other appetite suppressants. Consistent with prior research, the serotonergic precursor 5-HTP did not elicit head-weaving stereotypy.[13] Further analysis, employing network analysis and Markov transition matrices, revealed distinct behavioral transition probabilities associated with head-weaving. This stereotypy emerged as a dominant attractor state under the influence of certain drugs, highlighting subtle differences in their potential mechanisms of action. Indeed, this analytical approach demonstrates that head-weaving stereotypy was more likely to manifest as an ‘attractor’ behavior when dopaminergic pathways were modulated. This observation underscores the role of dopamine reuptake inhibitors – specifically phentermine, mazindol, diethylpropion, and cathine – in precipitating this repetitive motor behavior. Finally, sex differences were observed, primarily for tesofensine and d-amphetamine, suggesting a potential sexually dimorphic effect on stereotypy expression.

Hierarchical clustering and t-distributed Stochastic Neighbor Embedding (t-SNE) analyses further elucidating the distinct motor profiles elicited by the investigated appetite suppressants. Specifically, animals administered drugs primarily targeting serotonergic pathways – namely, 5-HTP– clustered closely with control groups. This proximity reflects behavioral profiles characterized by lower levels of stereotypy and more balanced transitions between quiet-awake, grooming, and locomotor states. In contrast, drugs modulating dopamine reuptake formed a distinct cluster, marked by pronounced stereotypy and locomotor behaviors, indicative of a characteristic dopaminergic influence on motor control. Interestingly, female rats treated with tesofensine formed an intermediary cluster, suggesting a partial overlap in motor profiles with both control and stereotypy-inducing drug groups. This intermediate positioning may reflect a mixed mechanism of action for tesofensine in females, potentially involving both serotonergic and subtle dopaminergic effects, or alternatively, enhanced sensitivity to the latter.

### Sexually dimorphic responses to appetite suppressants: Implications for weight loss and stereotypy

A key finding of this investigation is the pronounced sexually dimorphic response to specific appetite suppressants, impacting both weight loss and the manifestation of stereotyped behaviors. Cathine administration produced a striking sex difference in weight reduction, inducing weight loss exclusively in male rats (**Fig 1**), although both sexes exhibited robust head-weaving stereotypy in response to the drug (**Fig 6A**). In contrast, diethylpropion and tesofensine elicited significantly greater weight loss in female rats compared to their male counterparts (**Fig 1**). Regarding stereotypy, diethylpropion induced pronounced stereotypy in both sexes, whereas tesofensine predominantly elicited this behavior in female rats. Future research should elucidate the molecular mechanisms underpinning these observed dimorphic responses to appetite suppressants. Given these drugs’ capacity to inhibit norepinephrine (NET), serotonin (SERT), and dopamine (DAT) transporters, it is plausible that these transporters exhibit differential potencies and/or expression levels between male and female rats.[67,68] Alternatively, gonadal hormones may modulate susceptibility to appetite suppressants, contributing to the observed sex differences. Consequently, these hypotheses warrant further in-depth investigation.

### The striatum as a potential neural substrate for stereotypy induced by appetite suppressants

The striatum, particularly via dopaminergic (DA) signaling, plays a well-established role in the expression of stereotyped behaviors. This is evidenced by studies using animal models of dopaminergic supersensitivity resulting from DA deficiency.[3,48,69] For instance, while L-3,4-Dihydroxyphenylalanine (L-DOPA) administration has no discernible effect on wild-type mice, it induces a behavioral shift in dopamine-deficient mice when striatal DA levels surpass 50% of those found in wild-type counterparts. This shift manifests as a transition from hyperlocomotion to intense, focused stereotypy, concomitant with the upregulation of c-fos mRNA expression in the dorsal and ventrolateral striatum.[44] Furthermore, D1 receptor antagonists attenuate both L-DOPA-induced stereotypies and c-fos expression, highlighting the critical involvement of D1 receptor-expressing medium spiny neurons (MSNs) in mediating these stereotyped behaviors. Hypersensitivity to DA, as seen in these models, predisposes individuals to stereotypy upon hyperactivation of dopaminergic circuits, which can also be triggered by chronic exposure to psychomotor stimulants like cocaine and d-amphetamine.[70] Indeed, the striatum’s essential role in expressing motor stereotypies via dopaminergic mechanisms has been consistently demonstrated.[33,61,69] Moreover, it has been shown that dendritic atrophy and dendritic spine loss in dorsal striatum D1-MSNs from mice with repetitive behavior.[71] As shown here, the induction of robust motor stereotypies by appetite suppressants, coupled with their ability to curb hunger, further underscores the shared reliance of both appetite and motor circuits on DA as a key neuromodulator.

In this regard and beyond motor effects, DA efflux on the dorsal striatum has also been related to food intake and food reward,[72] in particular, caloric detection regardless of taste,[40] substantiating the idea that overlapping neural circuits are shared between caloric detection and motor effects, whereas DA in the ventral striatum, including the nucleus accumbens is related to both calories and taste hedonics of food.[40,73] In this regard, we have recently discovered that phentermine, diethylpropion, and cathine induced their appetite-suppressant properties and motor effects via DA D1 and D2 receptors in the nucleus accumbens,[10,42] highlighting the importance of DA in ventral and dorsal striatum on feeding and motor stereotypies.

Beyond the connection between food intake and motor circuits, further evidence points to a shared role of dopamine function in both obesity and drug addiction. Individuals with obesity and those with drug addiction exhibit similar impairments in dopaminergic circuits.[74] This aligns with the well-established role of some amphetamine derivatives as appetite suppressants. These drugs, by inhibiting the DAT transporter, increase dopamine levels, and thus, they suppress appetite and promote weight loss, but their addictive potential and motor side effects could also limit their use for the long-term treatment of obesity.

### Other types of motor stereotypies

Informal video analysis identified a novel stereotypy characterized by backward locomotion, which we termed “moonwalk stereotypy.” Notably, backward locomotion is known to be induced by hallucinogenic drugs such as lysergic acid diethylamide (LSD), a potent agonist at the serotoninergic 5-HT2a receptor.[32,75] Therefore, it is plausible that diethylpropion, beyond its dopaminergic effects, may also activate 5-HT2a receptors, potentially explaining the sporadic induction of backward locomotion observed. These findings are consistent with existing literature on amphetamine-induced psychosis, which can accelerate psychosis onset in susceptible individuals.[76,77] Furthermore, this may offer a potential explanation for rare case reports, primarily in women, of psychosis and hallucinations associated with the use of phentermine,[78,79] diethylpropion,[78,80–82] and mazindol.[83]

An important advantage of video recording using a bottom view was the discovery that during head-weaving stereotypy, rats also stuck out their tongue (tongue protrusion and retraction in the air). It has been reported that apomorphine, a potent dopaminergic agonist, more clearly induces this type of orolingual stereotypy,[61] but amphetamine and cocaine also induced this behavior, although to a lesser extent.[30,61] Here, we observed this behavior as particularly strong after administering mazindol (see **S1 Video** and **S2 Video**). Likewise, we have recently reported that a very high dose of tesofensine 6 mg/kg is needed to induce this orolingual dyskinesia in male rats,[25] and here we also found it in female rats treated with 2 mg/Kg. Recent optogenetic experiments have found that optogenetic activation of the projections between GABAergic neurons in the lateral hypothalamus to GABA neurons in VTA can induce food intake but also a strong gnawing and aberrant licking in the air or the floor stereotypy.[30]

### Appetite suppressants and spontaneous ejaculations

Unexpectedly, some appetite suppressants induced spontaneous ejaculations. Specifically, this effect was observed following administration of phentermine, mazindol, diethylpropion, and cathine. In contrast, d-amphetamine (primarily dopaminergic) and 5-HTP (primarily serotonergic) did not induce semen discharge in rats. This suggests that the observed ejaculations may be attributable to the combined effects of these drugs on both dopamine and serotonin signaling in brain regions involved in sexual function,[62–64] like the nucleus accumbens.[84,85] This interpretation is supported by reports of synergistic interactions between dopamine and serotonin on semen discharge.[64–66]

### Limitations of the study

We acknowledge that our video recordings did not allow us to systematically quantify orolingual dyskinesia, moonwalk stereotypy, or other more complex forms of stereotypy. Similarly, our method did not differentiate immobility from sleep, categorizing both as a “quiet-awake” state. Moreover, other classic forms of stereotypy induced by amphetamine, like “taffy pulling’’ a continuous movement of clasped forepaws toward and then away from the mouth, were not easily detected by our algorithm.[86] Further research should refine the automated ethogram by incorporating methods for precise detection and quantification of complex stereotypies and sleep behaviors. We anticipate that this open-source database[87] could facilitate and improve the study of drug-induced stereotypy in rats.

## Conclusion

In summary, we conclude that stereotypy behavior is not a homogeneous or static behavioral state but rather a broad spectrum and rich choreography of diverse motor acts that constantly alternate and compete for expression.[70] Crucially, appetite suppressants may exhibit sexually dimorphic responses, affecting males and females differently in terms of both weight loss and motor stereotypy effects. This underscores the importance of considering sex as a critical factor in their prescription and personalized use to optimize efficacy and minimize potential side effects. Indeed, our findings demonstrate that females exhibit greater sensitivity not only to the therapeutic effects but also to the adverse side effects of these compounds. This sex-dependent sensitivity to weight loss drugs extends beyond the classical appetite suppressants described here, encompassing newer generations of anti-obesity medications with different mechanisms of action. For example, in humans, females experience greater average weight loss than males when treated with the GLP-1 receptor agonist semaglutide (Ozempic), even at the same dose.[88,89] This observation suggests a broader role for sex hormones in modulating the physiological response to weight loss interventions, extending beyond appetite modulation.

## Data Availability Statement

The database and DeepLabCut trained network have been uploaded to the Open Science Framework[87] and are accessible via DOI: 10.17605/OSF.IO/MVD2U

## Acknowledgments

All behavioral task schemes were created using BioRender. We thank Fabiola Olvera Hernandez Mario Gil Moreno and Carlos Giovanni Sam Miranda for their valuable assistance with animal care.

## Competing Interests

The authors have declared that no competing interests exist.

## Financial Disclosure

This project was supported in part by CONAHCyT CF-2023-G-518 to R.G. The funders had no role in study design, data collection, and analysis, the decision to publish, or the preparation of the manuscript.

## Supplementary Figures

**S1 Fig.**
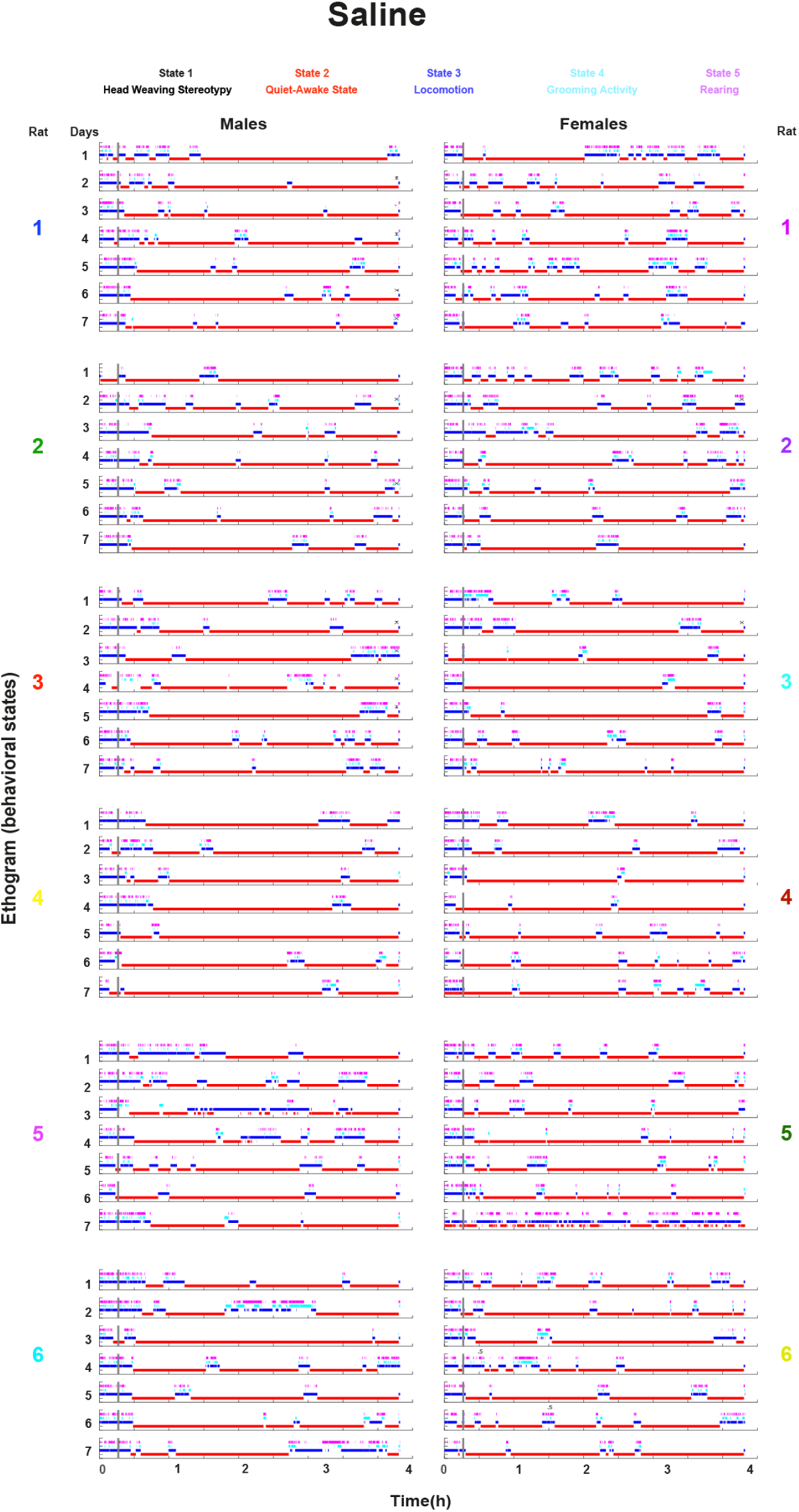
Automatic ethograms revealed that saline-treated rats exhibited no head-weaving stereotypy. Video frames were analyzed to classify rat behavior over time. Each frame was assigned to one of five mutually exclusive behavioral categories: stereotypy, quiet-awake state, locomotion, grooming, or rearing. For visualization purposes, each behavior was represented by a distinct color: **Stereotypy:** Black, **Quiet-awake state:** Red, **Locomotion:** Blue, **Grooming:** Cyan, and **Rearing:** Magenta. The left column displays six panels, each representing one of the six male rats, showing the ethogram for each of the seven treatment days. The right column mirrors the left column but for the female rats.

**S2 Fig.**
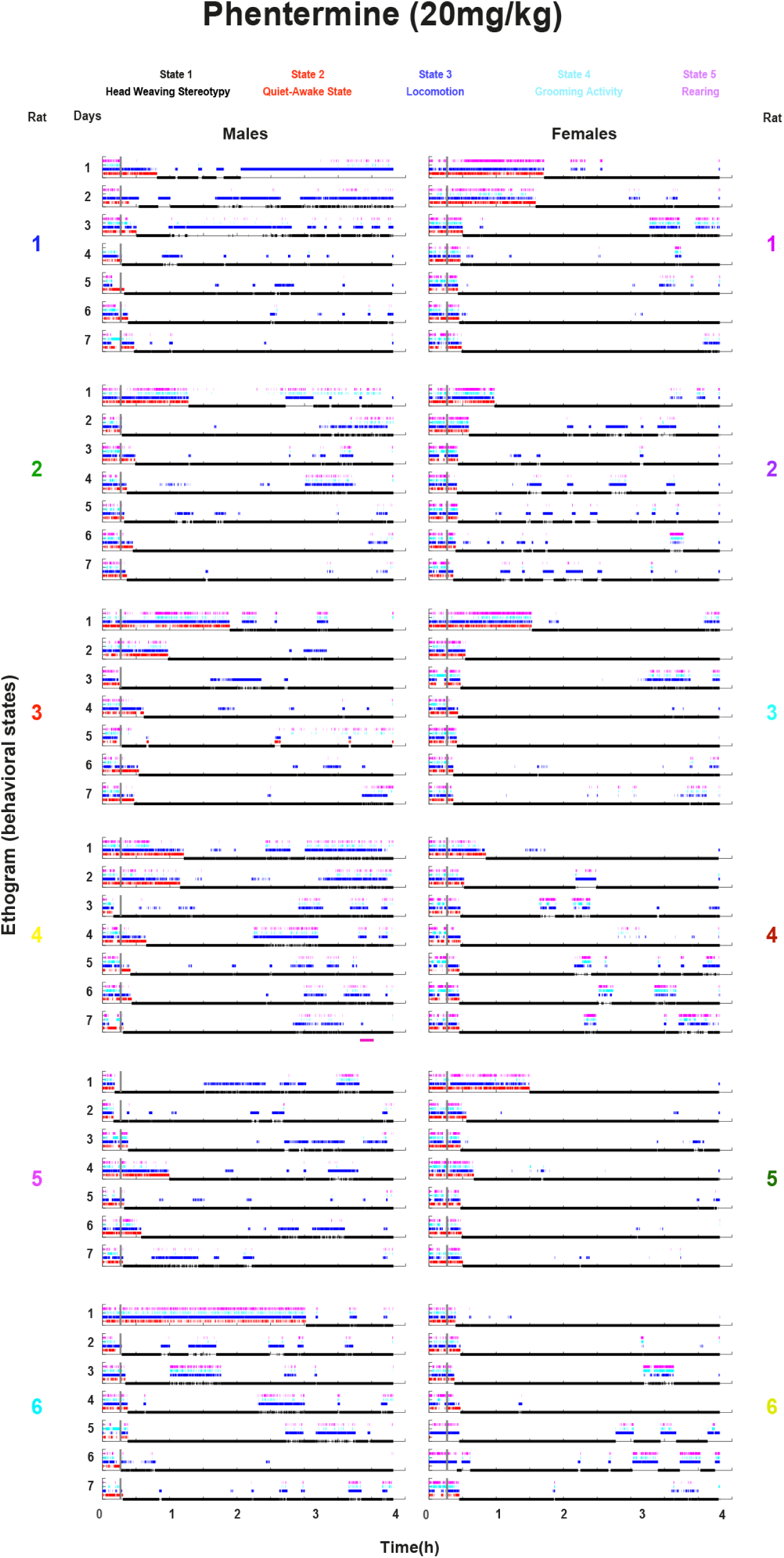
Automatic ethogram revealed that phentermine-treated rats exhibit head-weaving stereotypy that increases over the days. Same as for S. Fig 1. For visualization purposes, each behavior was represented by a distinct color: **Stereotypy:** Black, **Quiet-awake state:** Red, **Locomotion:** Blue, **Grooming:** Cyan, and **Rearing:** Magenta. The left column displays six panels, each representing one of the six male rats, showing the ethogram for each of the seven days of treatment. The right column mirrors the left column but for the female rats.

**S3 Fig.**
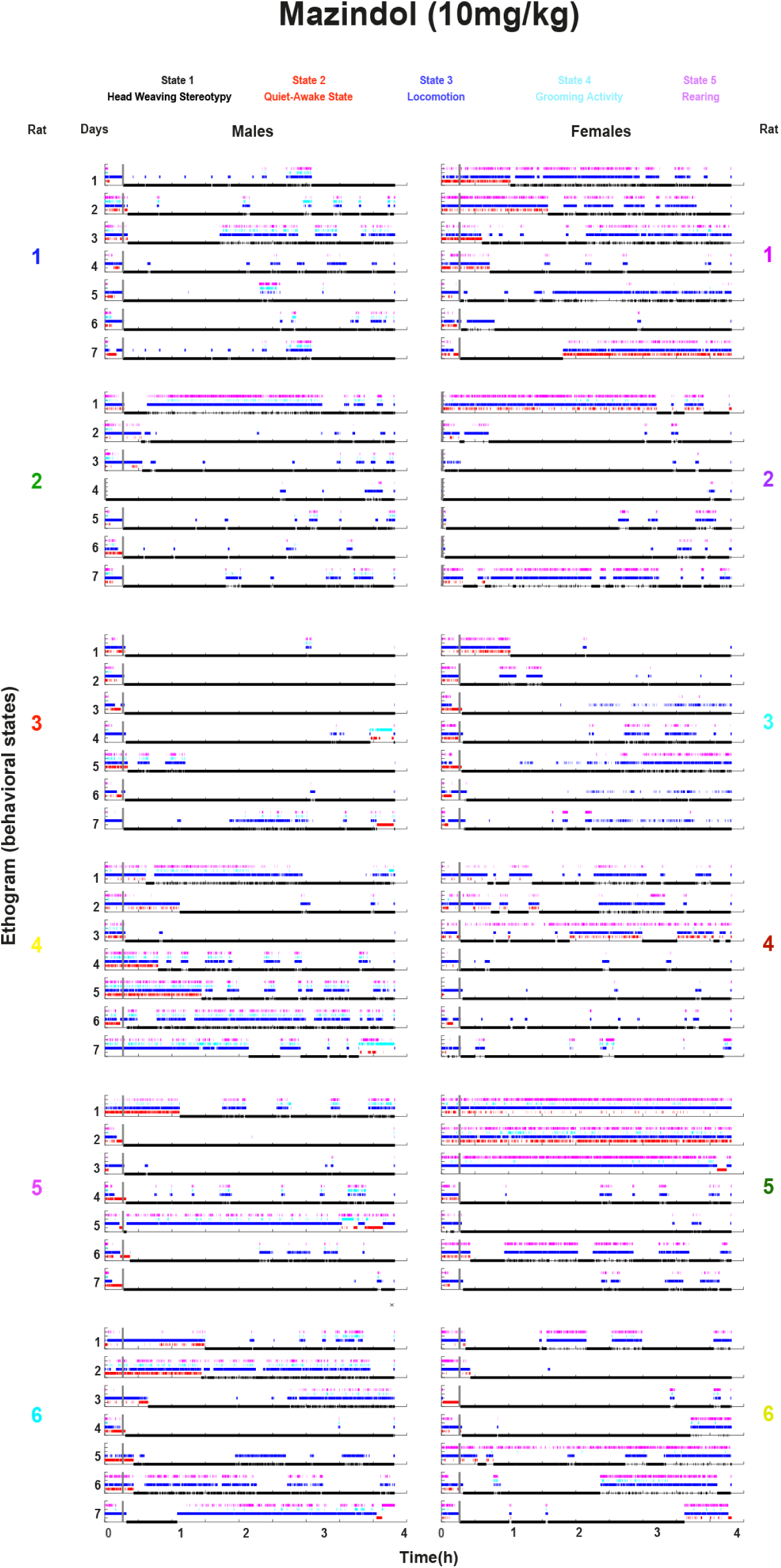
Automatic ethogram revealed that mazindol-treated rats exhibited head-weaving stereotypy similar in both females and males. Same as for **S1 Fig**. For visualization purposes, each behavior was represented by a distinct color: **Stereotypy:** Black, **Quiet-awake state:** Red, **Locomotion:** Blue, **Grooming:** Cyan, and **Rearing:** Magenta. The left column displays six panels, each representing one of the six male rats, showing the ethogram for each of the seven days of treatment. The right column mirrors the left column but for the female rats.

**S4 Fig.**
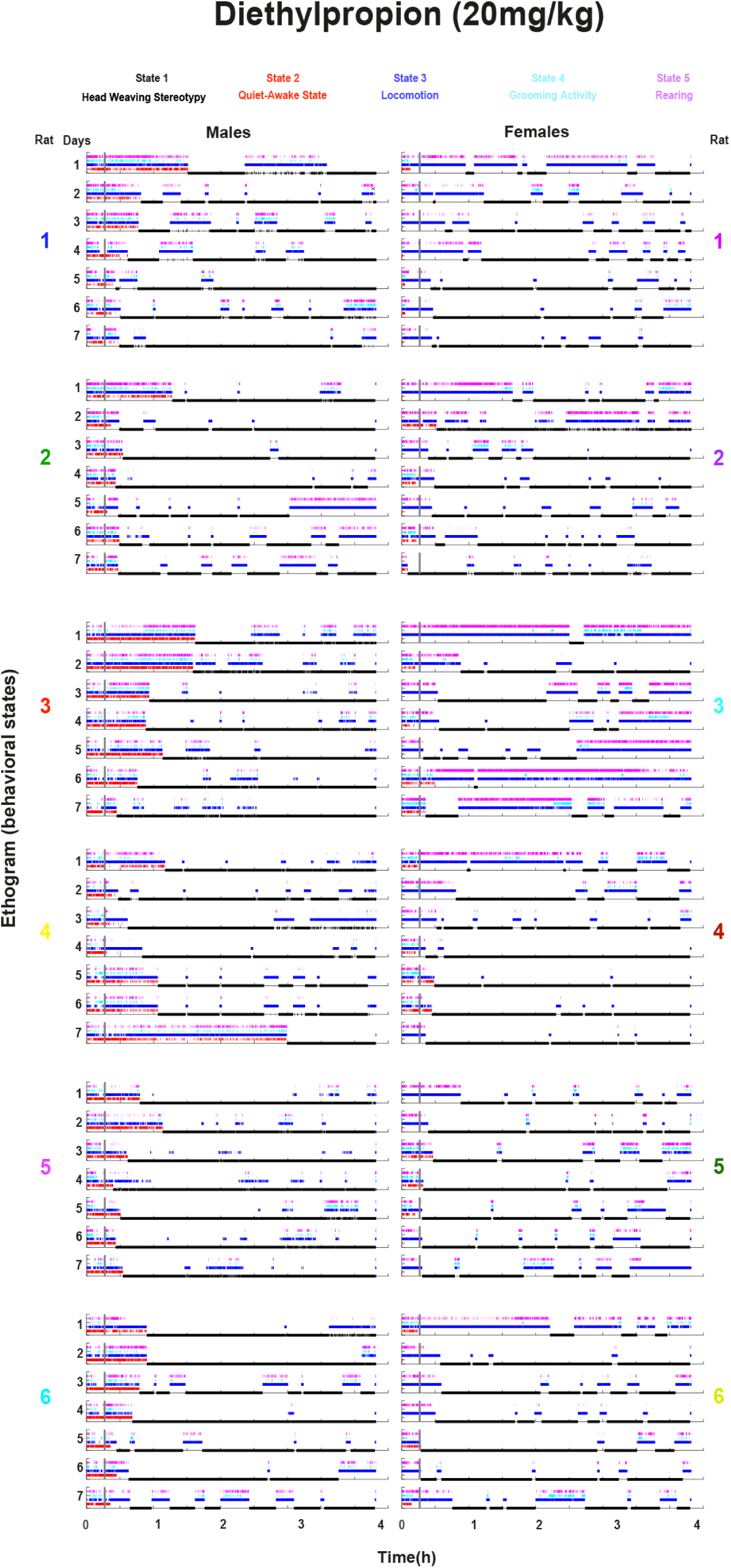
Automatic ethogram revealed that diethylpropion-treated rats exhibited a robust head-weaving stereotypy. Same as for **S1 Fig**. For visualization purposes, each behavior was represented by a distinct color: **Stereotypy:** Black, **Quiet-awake state:** Red, **Locomotion:** Blue, **Grooming:** Cyan, and **Rearing:** Magenta. The left column displays six panels, each representing one of the six male rats, showing the ethogram for each of the seven days of treatment. The right column mirrors the left column but for the female rats.

**S5 Fig.**
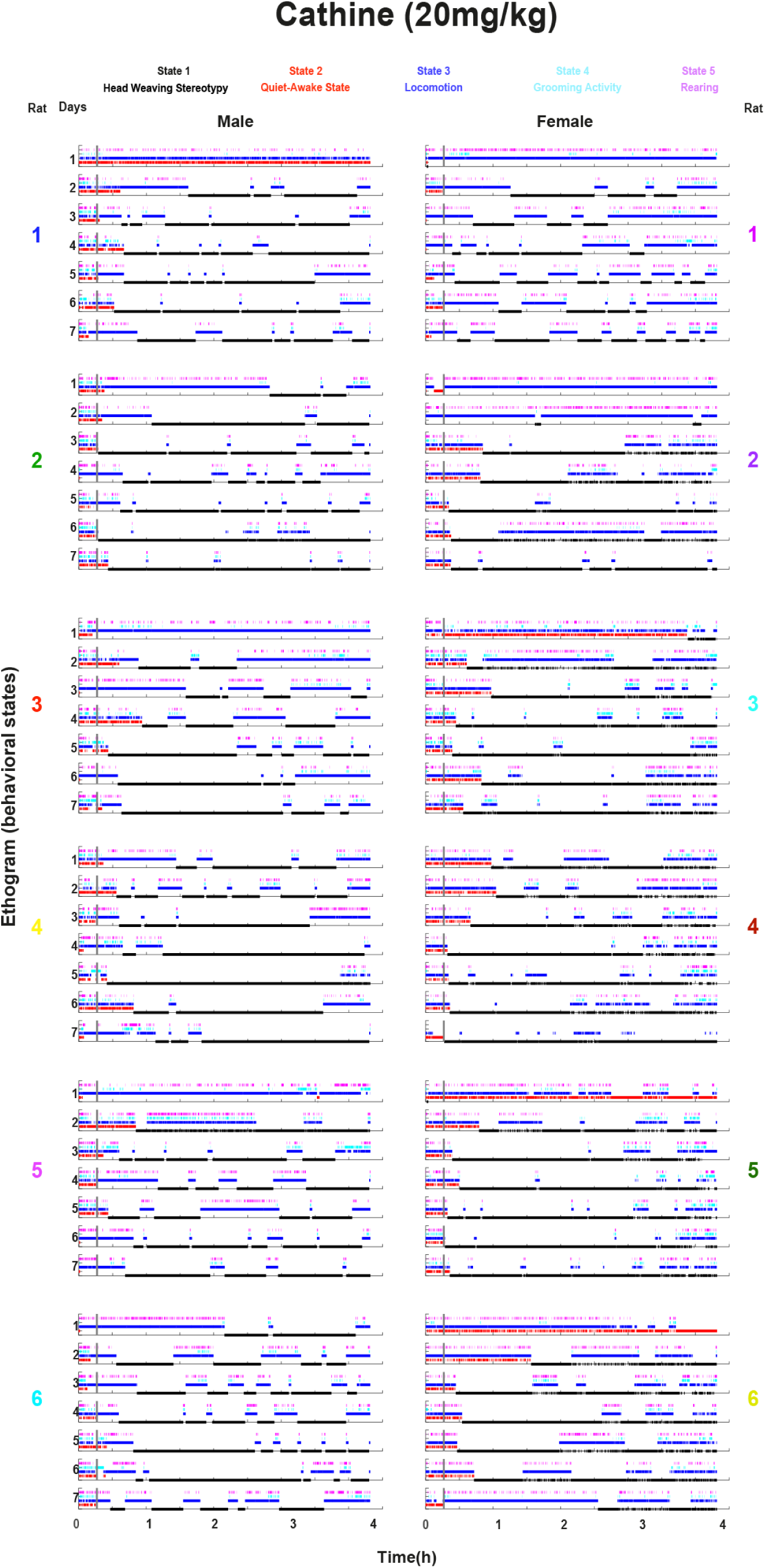
Automatic ethogram revealed that cathine-treated rats exhibited a heterogeneous head-weaving stereotypy. Same as for **S1 Fig**. For visualization purposes, each behavior was represented by a distinct color: **Stereotypy:** Black, **Quiet-awake state:** Red, **Locomotion:** Blue, **Grooming:** Cyan, and **Rearing:** Magenta. The left column displays six panels, each representing one of the six male rats, showing the ethogram for each of the seven days of treatment. The right column mirrors the left column but for the female rats.

**S6 Fig.**
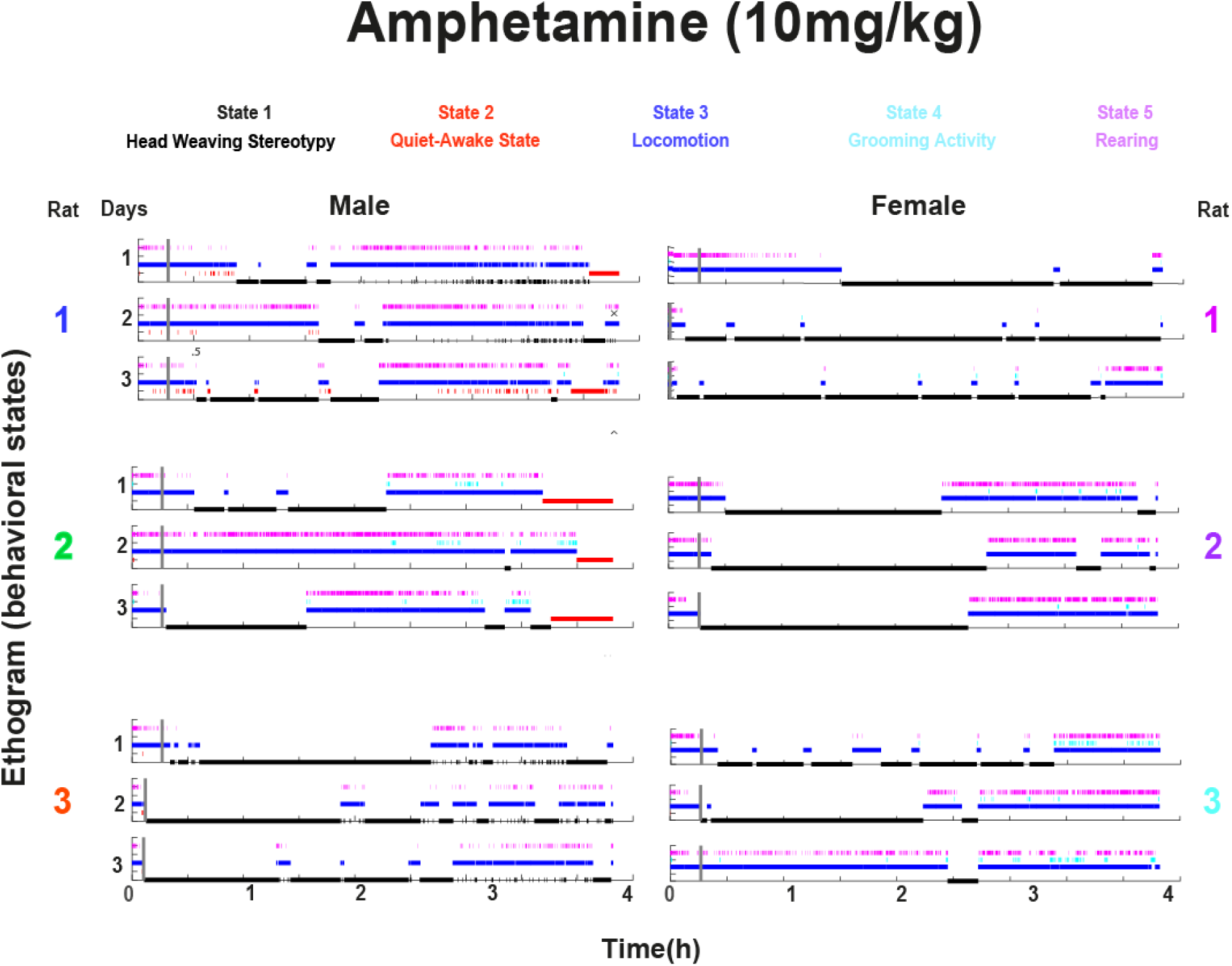
Automatic ethograms revealed that d-amphetamine-treated rats exhibited hyperlocomotion during the last hour of recording in females, while in males, it induced a quiet-awake state. Same as for **S1 Fig**. For visualization purposes, each behavior was represented by a distinct color: **Stereotypy:** Black, **Quiet-awake state:** Red, **Locomotion:** Blue, **Grooming:** Cyan, and **Rearing:** Magenta. The left column displays three panels, each representing one of the three male rats, showing the ethogram for each of the three days of treatment. The right column mirrors the left column but for the female rats.

**S7 Fig.**
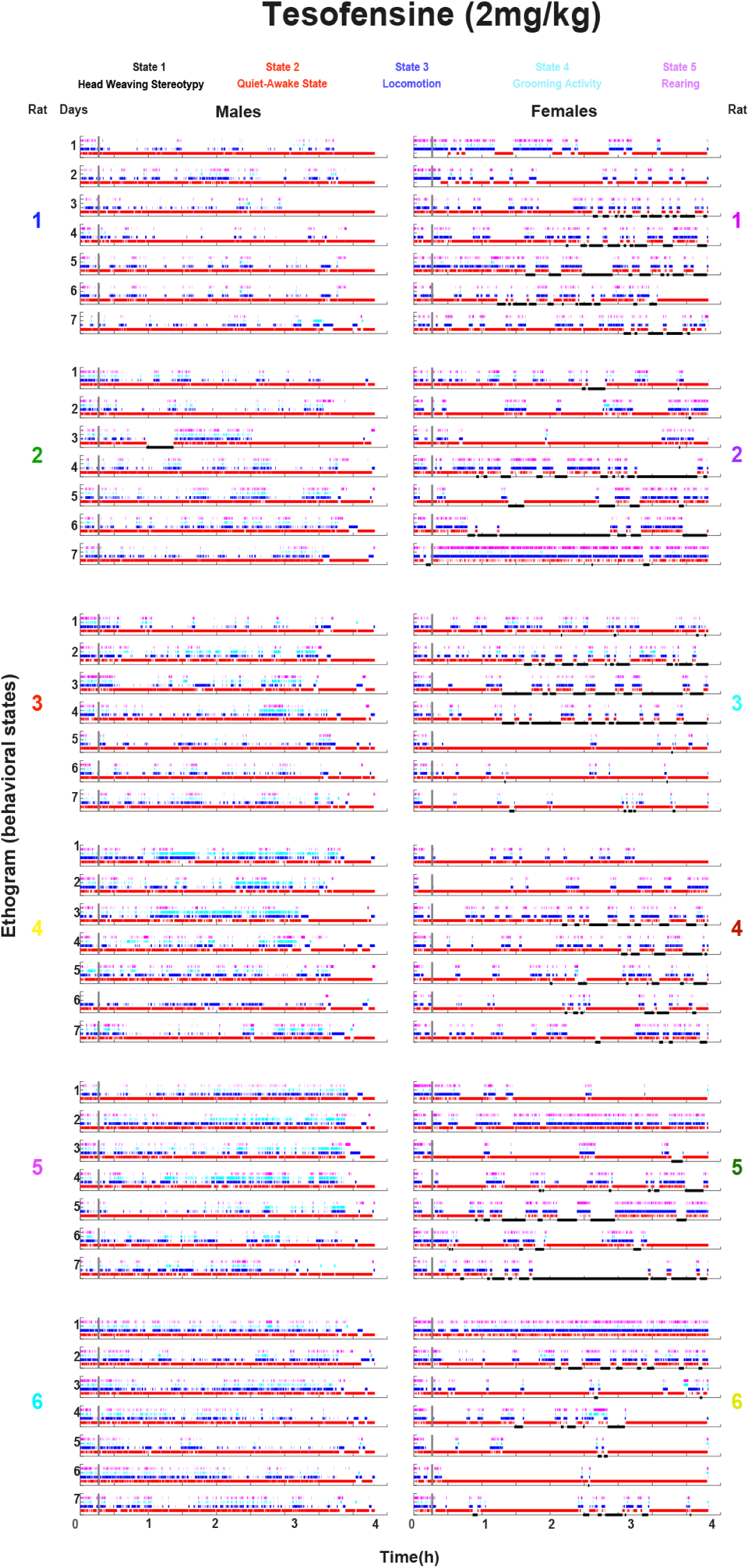
Automatic ethograms revealed that tesofensine-treated female rats exhibited head-weaving stereotypy, while in males, it induced a quiet-awake state. Same as for **S1 Fig**. For visualization purposes, each behavior was represented by a distinct color: **Stereotypy:** Black, **Quiet-awake state:** Red, **Locomotion:** Blue, **Grooming:** Cyan, and **Rearing:** Magenta. The left column displays six panels, each representing one of the six male rats, showing the ethogram for each of the seven days of treatment. The right column mirrors the left column but for the female rats.

**S8 Fig.**
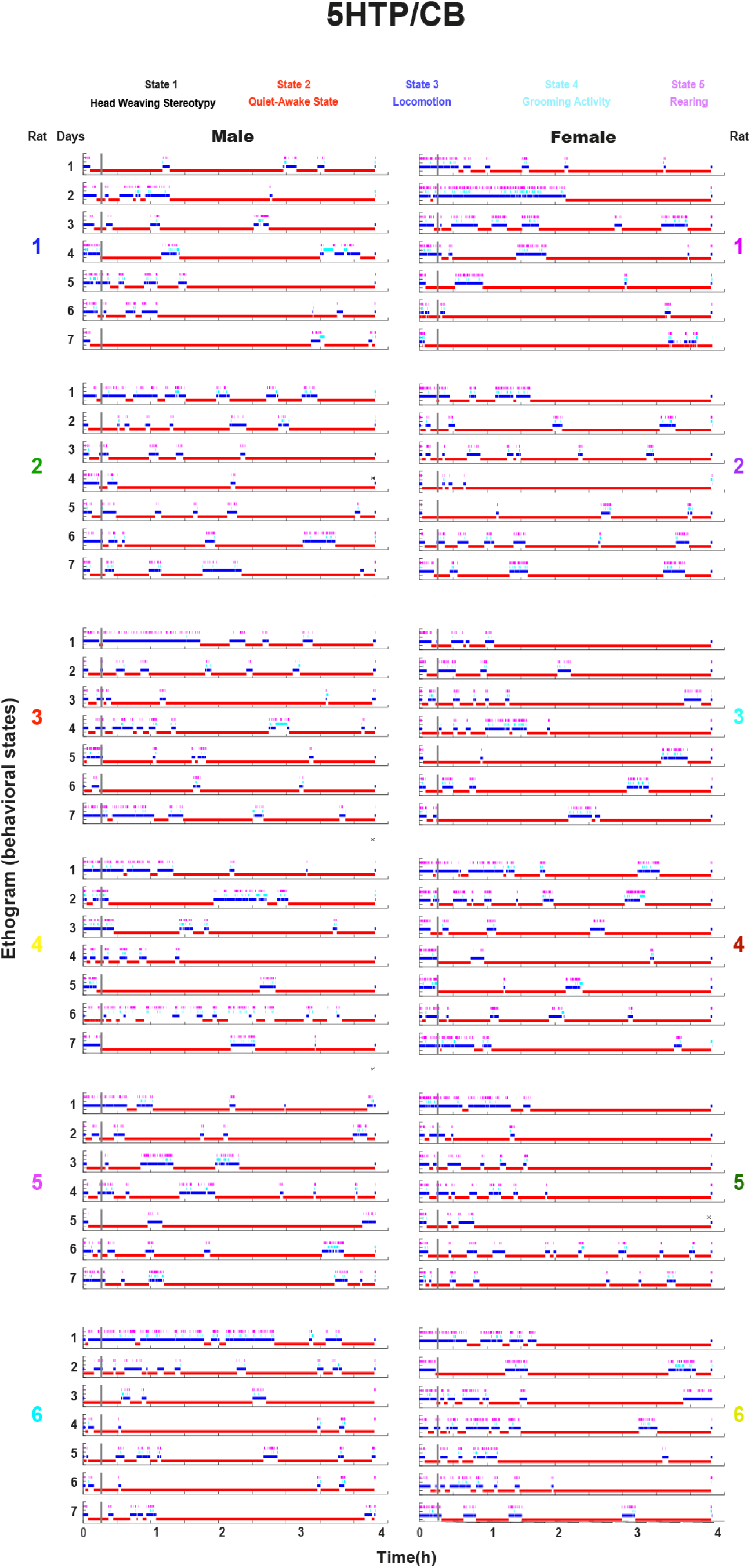
Automatic ethograms revealed that 5-HTP-treated rats exhibited no head-weaving stereotypy. Same as for **S1 Fig**. For visualization purposes, each behavior was represented by a distinct color: **Stereotypy:** Black, **Quiet-awake state:** Red, **Locomotion:** Blue, **Grooming:** Cyan, and **Rearing:** Magenta. The left column displays six panels, each representing one of the six male rats, showing the ethogram for each of the seven days of treatment. The right column mirrors the left column but for the female rats.

## Tables

S1 Table Statistical analysis for **Figure 1**

S2 Table Statistical analysis for **Figures 3 and 4**

S3 Table Time spent in each behavioral state

## Videos

S1 Video Head Weaving Stereotypy Males

S2 Video Head Weaving Stereotypy Females

S3 Video Locomotion Saline vs Amphetamine 10 mg/kg

S4 Video Orolingual Dyskinesia_Tesofensine 2 mg/kg

S5 Video Moonwalking Stereotypy_Male_Phentermine 20 mg/kg

S6 Video Moonwalking Stereotypy_Male_Diethylpropion 20 mg/kg

S7 Video Moonwalking Stereotypy_Female_Diethylpropion 20 mg/kg

S8 Video Moonwalking Stereotypy_Male_Mazindol 10 mg/kg

S9 Video Moonwalking Stereotypy_Female_Mazindol 10 mg/kg

S10 Video Moonwalking Stereotypy_Female_Amphetamine 20 mg/kg

S11 Video Spontaneous ejaculation_Phentermine 20 mg/kg

S12 Video Spontaneous ejaculation_Mazindol 20 mg/kg

S13 Video Spontaneous ejaculation_Diethylpropion 20 mg/kg

S14 Video Spontaneous ejaculation_Cathine 20 mg/kg

## Notes

https://osf.io/mvd2u/

### Competing Interest Statement

The authors have declared no competing interest.

## References

1. Papavramidou N, Christopoulou-Aletra H. Management of obesity in the writings of Soranus of Ephesus and Caelius Aurelianus. Obes Surg. 2008;18: 763–765. doi:10.1007/s11695-007-9362-1

2. Silverstone T. Appetite suppressants. A review. Drugs. 1992;43: 820–836. doi:10.2165/00003495-199243060-00003

3. Twohig M, Varra E. Treatment of Drug-Induced Stereotypy. Behav Anal Today. 2006;7. doi:10.1037/h0100081

4. Joo JK, Lee KS. Pharmacotherapy for obesity. J Menopausal Med. 2014;20: 90–96. doi:10.6118/jmm.2014.20.3.90

5. Valenza M, Steardo L, Cottone P, Sabino V. Diet-induced obesity and diet-resistant rats: differences in the rewarding and anorectic effects of d-amphetamine. Psychopharmacology (Berl). 2015;232: 3215–3226. doi:10.1007/s00213-015-3981-3

6. Tatsuta T, Kitanaka N, Kitanaka J, Morita Y, Takemura M. Effects of monoamine oxidase inhibitors on methamphetamine-induced stereotypy in mice and rats. Neurochem Res. 2005;30: 1377–1385. doi:10.1007/s11064-005-8390-2

7. Berman SM, Kuczenski R, McCracken JT, London ED. Potential Adverse Effects of Amphetamine Treatment on Brain and Behavior: A Review. Mol Psychiatry. 2009;14: 123–142. doi:10.1038/mp.2008.90

8. Khoramizadeh M, Effatpanah M, Mostaghimi A, Rezaei M, Mahjoub A, Shishehgar S. Treatment of amphetamine abuse/use disorder: a systematic review of a recent health concern. Daru J Fac Pharm Tehran Univ Med Sci. 2019;27: 743– 753. doi:10.1007/s40199-019-00282-3

9. LiverTox: Clinical and Research Information on Drug-Induced Liver Injury. Bethesda (MD): National Institute of Diabetes and Digestive and Kidney Diseases; 2012. Available: http://www.ncbi.nlm.nih.gov/books/NBK547852/

10. Kalyanasundar B, Solorio J, Perez CI, Hoyo-Vadillo C, Simon SA, Gutierrez R. The efficacy of the appetite suppressant, diethylpropion, is dependent on both when it is given (day vs. night) and under conditions of high fat dietary restriction. Appetite. 2016;100: 152–161. doi:10.1016/j.appet.2016.01.036

11. Hendricks EJ, Rothman RB, Greenway FL. How physician obesity specialists use drugs to treat obesity. Obes Silver Spring Md. 2009;17: 1730–1735. doi:10.1038/oby.2009.69

12. Hendricks EJ, Greenway FL, Westman EC, Gupta AK. Blood pressure and heart rate effects, weight loss and maintenance during long-term phentermine pharmacotherapy for obesity. Obes Silver Spring Md. 2011;19: 2351–2360. doi:10.1038/oby.2011.94

13. Perez CI, Kalyanasundar B, Moreno MG, Gutierrez R. The Triple Combination Phentermine Plus 5-HTP/Carbidopa Leads to Greater Weight Loss, With Fewer Psychomotor Side Effects Than Each Drug Alone. Front Pharmacol. 2019;10: 1327. doi:10.3389/fphar.2019.01327

14. Kalix P. A comparison of the catecholamine releasing effect of the khat alkaloids (-)-cathinone and (+)-norpseudoephedrine. Drug Alcohol Depend. 1983;11: 395–401. doi:10.1016/0376-8716(83)90031-5

15. Kalyanasundar B, Perez CI, Arroyo B, Moreno MG, Gutierrez R. The Appetite Suppressant D-norpseudoephedrine (Cathine) Acts via D1/D2-Like Dopamine Receptors in the Nucleus Accumbens Shell. Front Neurosci. 2020;14: 572328. doi:10.3389/fnins.2020.572328

16. Eddy NB, Halbach H, Isbell H, Seevers MH. Drug dependence: its significance and characteristics. Bull World Health Organ. 1965;32: 721–733.

17. Zelger JL, Schorno HX, Carlini EA. Behavioural effects of cathinone, an amine obtained from Catha edulis Forsk.: comparisons with amphetamine, norpseudoephedrine, apomorphine and nomifensine. Bull Narc. 1980;32: 67–81.

18. Ms E, Tj M, Hi S. A comparison of the effects of phenylpropanolamine, d-amphetamine and d-norpseudoephedrine on open-field locomotion and food intake in the rat. Appetite. 1987;9. doi:10.1016/0195-6663(87)90051-1

19. Neumann J, Hußler W, Hofmann B, Gergs U. Contractile Effects of Amphetamine, Pseudoephedrine, Nor-pseudoephedrine (Cathine), and Cathinone on Atrial Preparations of Mice and Humans. J Cardiovasc Pharmacol. 2024;83: 243–250. doi:10.1097/FJC.0000000000001536

20. Murphy JE, Donald JF, Molla AL, Crowder D. A comparison of mazindol (Teronac) with diethylpropion in the treatment of exogenous obesity. J Int Med Res. 1975;3: 202–206. doi:10.1177/030006057500300310

21. Sikdar SK, Oomura Y, Inokuchi A. Effects of mazindol on rat lateral hypothalamic neurons. Brain Res Bull. 1985;15: 33–38. doi:10.1016/0361-9230(85)90058-9

22. Wellman PJ. Systemic mazindol reduces food intake in rats via suppression of meal size and meal number. J Psychopharmacol Oxf Engl. 2008;22: 532–535. doi:10.1177/0269881107083837

23. Astrup A, Meier DH, Mikkelsen BO, Villumsen JS, Larsen TM. Weight Loss Produced by Tesofensine in Patients With Parkinson’s or Alzheimer’s Disease. Obesity. 2008;16: 1363–1369. doi:10.1038/oby.2008.56

24. Astrup A, Madsbad S, Breum L, Jensen TJ, Kroustrup JP, Larsen TM. Effect of tesofensine on bodyweight loss, body composition, and quality of life in obese patients: a randomised, double-blind, placebo-controlled trial. The Lancet. 2008;372: 1906–1913. doi:10.1016/S0140-6736(08)61525-1

25. Perez CI, Luis-Islas J, Lopez A, Diaz X, Molina O, Arroyo B, et al. Tesofensine, a novel antiobesity drug, silences GABAergic hypothalamic neurons. PloS One. 2024;19: e0300544. doi:10.1371/journal.pone.0300544

26. van Galen KA, ter Horst KW, Serlie MJ. Serotonin, food intake, and obesity. Obes Rev. 2021;22: e13210. doi:10.1111/obr.13210

27. Fog R. Stereotyped and non-stereotyped behaviour in rats induced by various stimulant drugs. Psychopharmacologia. 1969;14: 299–304. doi:10.1007/BF02190114

28. Fibiger HC, Fibiger HP, Zis AP. Attenuation of amphetamine-induced motor stimulation and stereotypy by 6-hydroxydopamine in the rat. Br J Pharmacol. 1973;47: 683–692. doi:10.1111/j.1476-5381.1973.tb08194.x

29. Creese I, Iversen SD. The pharmacological and anatomical substrates of the amphetamine response in the rat. Brain Res. 1975;83: 419–436. doi:10.1016/0006-8993(75)90834-3

30. Nieh EH, Matthews GA, Allsop SA, Presbrey KN, Leppla CA, Wichmann R, et al. Decoding neural circuits that control compulsive sucrose seeking. Cell. 2015;160: 528–541. doi:10.1016/j.cell.2015.01.003

31. Randrup A, Munkvad I. Pharmacology and physiology of stereotyped behavior. J Psychiatr Res. 1974;11: 1–10. doi:10.1016/0022-3956(74)90062-4

32. Yamamoto T, Ueki S. The role of central serotonergic mechanisms on head-twitch and backward locomotion induced by hallucinogenic drugs. Pharmacol Biochem Behav. 1981;14: 89–95. doi:10.1016/0091-3057(81)90108-8

33. Robbins TW, de Wied D. Mode of action of apomorphine and dexamphetamine on gnawing compulsion in rats: A.M. Ernst. Psychopharmacologia (Berl.) 10, 316–323 (1967). Psychopharmacology (Berl). 1991;103: 285–286. doi:10.1007/BF02244279

34. Balsara JJ, Jadhav JH, Muley MP, Chandorkar AG. Effect of drugs influencing central serotonergic mechanisms on methamphetamine-induced stereotyped behavior in the rat. Psychopharmacology (Berl). 1979;64: 303–307. doi:10.1007/BF00427514

35. Canales JJ, Gilmour G, Iversen SD. The role of nigral and thalamic output pathways in the expression of oral stereotypies induced by amphetamine injections into the striatum. Brain Res. 2000;856: 176–183. doi:10.1016/s0006-8993(99)02344-6

36. Snow V, Barry P, Fitterman N, Qaseem A, Weiss K, Clinical Efficacy Assessment Subcommittee of the American College of Physicians. Pharmacologic and surgical management of obesity in primary care: a clinical practice guideline from the American College of Physicians. Ann Intern Med. 2005;142: 525–531. doi:10.7326/0003-4819-142-7-200504050-00011

37. Cooke D, Bloom S. The obesity pipeline: current strategies in the development of anti-obesity drugs. Nat Rev Drug Discov. 2006;5: 919–931. doi:10.1038/nrd2136

38. Ioannides-Demos LL, Proietto J, Tonkin AM, McNeil JJ. Safety of drug therapies used for weight loss and treatment of obesity. Drug Saf. 2006;29: 277–302. doi:10.2165/00002018-200629040-00001

39. Ferreira JG, Tellez LA, Ren X, Yeckel CW, de Araujo IE. Regulation of fat intake in the absence of flavour signalling. J Physiol. 2012;590: 953–972. doi:10.1113/jphysiol.2011.218289

40. Tellez LA, Han W, Zhang X, Ferreira TL, Perez IO, Shammah-Lagnado SJ, et al. Separate circuitries encode the hedonic and nutritional values of sugar. Nat Neurosci. 2016;19: 465–470. doi:10.1038/nn.4224

41. de Lartigue G, McDougle M. Dorsal striatum dopamine oscillations: setting the pace of food anticipatory activity. Acta Physiol Oxf Engl. 2019;225: e13152. doi:10.1111/apha.13152

42. Kalyanasundar B, Perez CI, Luna A, Solorio J, Moreno MG, Elias D, et al. D1 and D2 antagonists reverse the effects of appetite suppressants on weight loss, food intake, locomotion, and rebalance spiking inhibition in the rat NAc shell. J Neurophysiol. 2015;114: 585–607. doi:10.1152/jn.00012.2015

43. Björklund A, Dunnett SB. Dopamine neuron systems in the brain: an update. Trends Neurosci. 2007;30: 194–202. doi:10.1016/j.tins.2007.03.006

44. Palmiter RD. Restricting Dopaminergic Signaling to Either Dorsolateral or Medial Striatum Facilitates Cognition. 20 Jan 2010 [cited 30 Sep 2024]. Available: https://www.jneurosci.org/content/jneuro/30/3/1158.full.pdf

45. Lobo MK, Nestler EJ. The striatal balancing act in drug addiction: distinct roles of direct and indirect pathway medium spiny neurons. Front Neuroanat. 2011;5: 41. doi:10.3389/fnana.2011.00041

46. Kelley AE. Measurement of Rodent Stereotyped Behavior. Curr Protoc Neurosci. 1998;4: 8.8.1–8.8.13. doi:10.1002/0471142301.ns0808s04

47. Am E. Mode of action of apomorphine and dexamphetamine on gnawing compulsion in rats. Psychopharmacologia. 1967;10. doi:10.1007/BF00403900

48. B C, Rj N. The role of telencephalic dopaminergic systems in the mediation of apomorphine-stereotyped behaviour. Eur J Pharmacol. 1973;24. doi:10.1016/0014-2999(73)90108-8

49. Beatty WW, Holzer GA. Sex differences in stereotyped behavior in the rat. Pharmacol Biochem Behav. 1978;9: 777–783. doi:10.1016/0091-3057(78)90356-8

50. Mathis A, Mamidanna P, Cury KM, Abe T, Murthy VN, Mathis MW, et al. DeepLabCut: markerless pose estimation of user-defined body parts with deep learning. Nat Neurosci. 2018;21: 1281–1289. doi:10.1038/s41593-018-0209-y

51. Guillory J. The Merck Index: An Encyclopedia of Chemicals, Drugs, and Biologicals Edited by Maryadele J. O’Neil, Patricia E. Heckelman, Cherie B. Koch, and Kristin J. Roman. Merck, John Wiley & Sons, Inc., Hoboken, NJ. 2006. xiv + 2564 pp. 18 × 26 cm. ISBN13 978-0-911910-001. $125.00. J Med Chem - J MED CHEM. 2007;50: 590–590. doi:10.1021/jm068049o

52. Pechnick R, Janowsky DS, Judd L. Differential effects of methylphenidate and d-amphetamine on stereotyped behavior in the rat. Psychopharmacology (Berl). 1979;65: 311–315. doi:10.1007/BF00492220

53. Arroyo B, Hernandez-Lemus E, Gutierrez R. The flow of reward information through neuronal ensembles in the accumbens. Cell Rep. 2024;43: 114838. doi:10.1016/j.celrep.2024.114838

54. Coss A, Suaste E, Gutierrez R. Lateral NAc Shell D1 and D2 Neuronal Ensembles Concurrently Predict Licking Behavior and Categorize Sucrose Concentrations in a Context-dependent Manner. Neuroscience. 2022;493: 81–98. doi:10.1016/j.neuroscience.2022.04.022

55. Baum LE, Petrie T, Soules G, Weiss N. A Maximization Technique Occurring in the Statistical Analysis of Probabilistic Functions of Markov Chains. Ann Math Stat. 1970;41: 164–171. doi:10.1214/aoms/1177697196

56. Robinson TE, Camp DM. Long-lasting effects of escalating doses of *d*-amphetamine on brain monoamines, amphetamine-induced stereotyped behavior and spontaneous nocturnal locomotion. Pharmacol Biochem Behav. 1987;26: 821–827. doi:10.1016/0091-3057(87)90616-2

57. James RC, Franklin MR. Tissue concentrations and the mortality of *d*-amphetamine in mice: Changes associated with the use of diethylaminoethyl 2,2-diphenylvalerate · HCl. Toxicol Appl Pharmacol. 1978;44: 75–80. doi:10.1016/0041-008X(78)90285-5

58. Fowler SC, Birkestrand B, Chen R, Vorontsova E, Zarcone T. Behavioral sensitization to amphetamine in rats: changes in the rhythm of head movements during focused stereotypies. Psychopharmacology (Berl). 2003;170: 167–177. doi:10.1007/s00213-003-1528-5

59. Fowler SC, Pinkston JW, Vorontsova E. Clozapine and prazosin slow the rhythm of head movements during focused stereotypy induced by d-amphetamine in rats. Psychopharmacology (Berl). 2007;192: 219–230. doi:10.1007/s00213-007-0705-3

60. Baumann MH, Williams Z, Zolkowska D, Rothman RB. Serotonin (5-HT) precursor loading with 5-hydroxy-l-tryptophan (5-HTP) reduces locomotor activation produced by (+)-amphetamine in the rat. Drug Alcohol Depend. 2011;114: 147–152. doi:10.1016/j.drugalcdep.2010.09.015

61. Canales JJ, Graybiel AM. A measure of striatal function predicts motor stereotypy. Nat Neurosci. 2000;3: 377–383. doi:10.1038/73949

62. Hsieh GC, Hollingsworth PR, Martino B, Chang R, Terranova MA, O’Neill AB, et al. Central mechanisms regulating penile erection in conscious rats: the dopaminergic systems related to the proerectile effect of apomorphine. J Pharmacol Exp Ther. 2004;308: 330–338. doi:10.1124/jpet.103.057455

63. Yonezawa A, Yoshizumi M, Ebiko M, Ise S, Watanabe C, Mizoguchi H, et al. Ejaculatory response induced by a 5-HT2 receptor agonist m-CPP in rats: differential roles of 5-HT2 receptor subtypes. Pharmacol Biochem Behav. 2008;88: 367–373. doi:10.1016/j.pbb.2007.09.009

64. Yonezawa A, Yoshizumi M, Ise S-N, Watanabe C, Mizoguchi H, Furukawa K, et al. Synergistic actions of apomorphine and m-chlorophenylpiperazine on ejaculation, but not penile erection in rats. Biomed Res Tokyo Jpn. 2009;30: 71–78. doi:10.2220/biomedres.30.71

65. Andersson KE. Pharmacology of erectile function and dysfunction. Urol Clin North Am. 2001;28: 233–247. doi:10.1016/s0094-0143(05)70134-8

66. Yoshizumi M, Yonezawa A, Kimura Y, Watanabe C, Sakurada S, Mizoguchi H. Central Mechanisms of Apomorphine and m-Chlorophenylpiperazine on Synergistic Action for Ejaculation in Rats. J Sex Med. 2021;18: 231–239. doi:10.1016/j.jsxm.2020.10.014

67. Gordon JH, Shellenberger MK. Regional catecholamine content in the rat brain: Sex differences and correlation with motor activity. Neuropharmacology. 1974;13: 129–137. doi:10.1016/0028-3908(74)90030-6

68. Costa KM, Schenkel D, Roeper J. Sex-dependent alterations in behavior, drug responses and dopamine transporter expression in heterozygous DAT-Cre mice. Sci Rep. 2021;11: 3334. doi:10.1038/s41598-021-82600-x

69. Chartoff EH, Marck BT, Matsumoto AM, Dorsa DM, Palmiter RD. Induction of stereotypy in dopamine-deficient mice requires striatal D1 receptor activation. Proc Natl Acad Sci U S A. 2001;98: 10451–10456. doi:10.1073/pnas.181356498

70. Pierce RC, Kalivas PW. A circuitry model of the expression of behavioral sensitization to amphetamine-like psychostimulants. Brain Res Brain Res Rev. 1997;25: 192–216. doi:10.1016/s0165-0173(97)00021-0

71. Engeln M, Song Y, Chandra R, La A, Fox ME, Evans B, et al. Individual differences in stereotypy and neuron subtype translatome with TrkB deletion. Mol Psychiatry. 2021;26: 1846–1859. doi:10.1038/s41380-020-0746-0

72. Han W, Tellez LA, Perkins MH, Perez IO, Qu T, Ferreira J, et al. A Neural Circuit for Gut-Induced Reward. Cell. 2018;175: 665–678.e23. doi:10.1016/j.cell.2018.08.049

73. de Araujo IE, Schatzker M, Small DM. Rethinking Food Reward. Annu Rev Psychol. 2020;71: 139–164. doi:10.1146/annurev-psych-122216-011643

74. Volkow ND, Wang G-J, Tomasi D, Baler RD. Obesity and addiction: neurobiological overlaps. Obes Rev. 2013;14: 2–18. doi:10.1111/j.1467-789X.2012.01031.x

75. Woolley DW. Production of abnormal (psychotic?) behavior in mice with lysergic acid diethylamide, and its partial prevention with cholinergic drugs and serotonin. Proc Natl Acad Sci. 1955;41: 338–344. doi:10.1073/pnas.41.6.338

76. Connell PH. Amphetamine Psychosis. Br Med J. 1957;1: 582.

77. Bramness JG, Gundersen ØH, Guterstam J, Rognli EB, Konstenius M, Løberg E-M, et al. Amphetamine-induced psychosis - a separate diagnostic entity or primary psychosis triggered in the vulnerable? BMC Psychiatry. 2012;12: 221. doi:10.1186/1471-244X-12-221

78. Hoffman BF. Diet pill psychosis. Can Med Assoc J. 1977;116: 351–355.

79. Cleare AJ. Phentermine, psychosis, and family history. J Clin Psychopharmacol. 1996;16: 470–471. doi:10.1097/00004714-199612000-00020

80. Petursson H. Diethylpropion and paranoid psychosis. Aust N Z J Psychiatry. 1979;13: 67–68. doi:10.3109/00048677909159112

81. Martin CA, Iwamoto ET. Diethylpropion-induced psychosis reprecipitated by an MAO inhibitor: case report. J Clin Psychiatry. 1984;45: 130–131.

82. Carney MW. Diethylpropion and psychosis. Clin Neuropharmacol. 1988;11: 183–188. doi:10.1097/00002826-198804000-00011

83. Willis JH. Letter: Abuse of non-amphetamine appetite suppressants. Lancet Lond Engl. 1976;1: 37. doi:10.1016/s0140-6736(76)92930-5

84. Fiorino DF, Phillips AG. Facilitation of sexual behavior and enhanced dopamine efflux in the nucleus accumbens of male rats after D-amphetamine-induced behavioral sensitization. J Neurosci Off J Soc Neurosci. 1999;19: 456–463. doi:10.1523/JNEUROSCI.19-01-00456.1999

85. Matsumoto J, Urakawa S, Hori E, de Araujo MFP, Sakuma Y, Ono T, et al. Neuronal responses in the nucleus accumbens shell during sexual behavior in male rats. J Neurosci Off J Soc Neurosci. 2012;32: 1672–1686. doi:10.1523/JNEUROSCI.5140-11.2012

86. Breese GR, Baumeister AA, McCown TJ, Emerick SG, Frye GD, Crotty K, et al. Behavioral differences between neonatal and adult 6-hydroxydopamine-treated rats to dopamine agonists: relevance to neurological symptoms in clinical syndromes with reduced brain dopamine. J Pharmacol Exp Ther. 1984;231: 343– 354.

87. Lopez A, Gutierrez R. Data base: Sex-specific effects of appetite suppressants and stereotypy in rats. 2025 [cited 4 Feb 2025]. doi:10.17605/OSF.IO/MVD2U

88. Rentzeperi E, Pegiou S, Koufakis T, Grammatiki M, Kotsa K. Sex Differences in Response to Treatment with Glucagon-like Peptide 1 Receptor Agonists: Opportunities for a Tailored Approach to Diabetes and Obesity Care. J Pers Med. 2022;12: 454. doi:10.3390/jpm12030454

89. Gasoyan H, Pfoh ER, Schulte R, Le P, Butsch WS, Rothberg MB. One-Year Weight Reduction With Semaglutide or Liraglutide in Clinical Practice. JAMA Netw Open. 2024;7: e2433326. doi:10.1001/jamanetworkopen.2024.33326

